# Revealing the functional traits that are linked to hidden environmental factors in community assembly

**DOI:** 10.1101/2020.09.10.292151

**Authors:** Valério D. Pillar, Francesco Maria Sabatini, Ute Jandt, Sergio Camiz, Helge Bruelheide

**Affiliations:** Department of Ecology, Universidade Federal do Rio Grande do Sul, Porto Alegre, RS, 91501-970, Brazil; Martin Luther University Halle-Wittenberg, Institute of Biology/Geobotany and Botanical Garden, Am Kirchtor 1, 06108 Halle, Germany; German Centre for Integrative Biodiversity Research (iDiv) Halle-Jena-Leipzig, Deutscher Platz 5e, 04103 Leipzig, Germany; National Research Council - ISPC, Piazzale Aldo Moro 5, 00185 Roma, Italy

**Keywords:** Beals smoothing, community assembly, environmental filtering, fuzzy-weighting, hidden environmental factors, species traits, species co-occurrence

## Abstract

**Aim:** To identify functional traits that best predict community assembly without knowing the driving environmental factors.

**Methods:** We propose a new method that is based on the correlation r(**XY**) between two matrices of potential community composition: matrix **X** is fuzzy-weighted by trait similarities of species, and matrix **Y** is derived by Beals smoothing using the probabilities of species co-occurrences. Since matrix **X** is based on one or more traits, r(**XY**) measures how well the traits used for fuzzy-weighting reflect the observed co-occurrence patterns. We developed an optimization algorithm that identifies those traits that maximize this correlation, together with an appropriate permutational test for significance. Using metacommunity data generated by a stochastic, individual-based, spatially explicit model, we assessed the type I error and the power of our method across different simulation scenarios, varying environmental filtering parameters, number of traits and trait correlation structures. We then applied the method to real-world community and trait data of dry calcareous grassland communities across Germany to identify, out of 49 traits, the combination of traits that maximizes r(**XY**).

**Results:** The method correctly identified the relevant traits involved in the community assembly mechanisms specified in simulations. It had high power and accurate type I error and was robust against confounding aspects related to interactions between environmental factors, strength of limiting factors, and correlation among traits. In the grassland dataset, the method identified five traits that best explained community assembly. These traits reflected the size and the leaf economics spectrum, which are related to succession and resource supply, factors that may not be always measured in real-world situations.

**Conclusions:** Our method successfully identified the relevant traits mediating community assembly driven by environmental factors which may be hidden for not being measured or accessible at the spatial or temporal scale of the study.

## Introduction

Understanding how species assemble in space and time is critical for predicting biodiversity responses to environmental factors (D’Amen et al. 2017) and the effects of biodiversity losses on ecosystem processes and services (Newbold 2018). In communities connected by dispersal, patterns of repeated co-occurrence and apparent mutual avoidance among species have often been observed (e.g. Diamond 1975; Münzbergová & Herben 2004). This is a consequence of the species’ ecological niches and interactions, both of which are mediated by species’ morphological, physiological, phenological, or behavioural characteristics, here collectively indicated as functional traits (Keddy 1992; McGill et al. 2006; Wilson 2007; Götzenberger et al. 2012). These “restrictions on the observed patterns” constitute community assembly rules (Wilson et al. 1999).

If community assembly is mediated by abiotic and biotic environmental factor-trait relations, species co-occurrence patterns may naturally arise, because species having similar traits will respond similarly to environmental factors. Imagine an environmental factor *e*_*1*_ affecting species performance via a trait *t*_*1*_, i.e., *e*_*1*_ -> *t*_*1*_. All else being equal, at a given level of *e*_*1*_ species will tend to co-occur with those having similar values of trait *t*_*1*_. This will generate trait convergence for *t_1_* or, in other words, a trend in community-weighted means (CWMs) along changing *e*_*1*_, i.e., *e*_*1*_ ->CWM_*t1*_. However, community assembly involves more complex mechanisms than that. First, the units subject to environmental filtering are whole organisms with sets of morpho-physio-phenological traits (Violle et al. 2007) which cannot be physically disentangled in response to different factors. Second, traits are often correlated, given that the multivariate trait space of species is strongly concentrated in a small number of trait value combinations, owing to coordination and trade-offs between traits as well as ecological and phylogenetic constraints (Murren 2002; Díaz et al. 2016; Céréghino et al. 2018). As a consequence of these two constraints, a factor effect (*e*_*1*_) on a trait (*t*_*1*_) may depend on the value of another trait (*t*_*2*_) in the same organism, either under the effect of the same factor, i.e., *e_1_* -> *t_1_*|*t_2_*, or another factor, i.e., (*e_1_* -> *t_1_*) | (*e_2_* -> *t_2_*). In this case, one trait may be more limiting than another depending on the strength of the factor effects (Sih & Gleeson 1995; Gorban et al. 2011). Also, unknown factors affecting *t_1_* will generate increased variance in *t*_*1*_ along the known *e*_*1*_ gradient (Kaiser et al. 1994; Thomson et al. 1996; Cade & Noon 2003). These mechanisms may generate patterns of trait divergence (Pillar et al. 2009), e.g., when the community-weighted variance, or functional diversity (FD), of a trait increases along an environmental gradient.

But how to identify which functional traits are relevant in mediating community assembly, irrespective of whether this depends on mechanisms leading to convergence or divergence patterns? Traditionally, these traits have been identified by relating community trait patterns to environmental conditions or resource levels, hereafter called *environmental factors* for simplicity (Pillar & Orlóci 1993; Díaz & Cabido 1997; Pillar 1999; Lavorel & Garnier 2002; Pillar et al. 2009; Bruelheide et al. 2018). This approach, however, falls short when these factors are hidden, i.e., unknown or not observable. This is the case, for instance, when the factor was simply not measured, when it is related to unknown past conditions, but also when it affects community assembly at a much finer resolution than the grain size of the studied community units. Moreover, community assembly might also depend on biotic factors related, for instance, to predation, competition, or facilitation. These factors are often difficult to measure, but are likewise expected to shape the functional profile of ecological communities (Mason & Wilson 2006; D’Amen et al. 2017).

Under the assumption that these relevant yet hidden factors are reflected in community composition, there might be a way for analysing compositional data which allows to highlight the fundamental traits mediating community assembly. Once the traits are known, one can use factor-trait relations known from ecological theory or from other empirical studies (e.g. Díaz et al. 2007; Dubuis et al. 2013; Bruelheide et al. 2018) to make inferences about the factors, even if hidden, which are responsible for filtering (Keddy 1992) species in the studied communities.

Here we propose and test a data-driven method to identify those functional traits that best predict community assembly without knowing the relevant environmental factors shaping the studied communities. The foundation of our approach is to relate two ways of predicting potential community composition to each other, either based on the probability of species co-occurrence (Beals, 1984) or using fuzzy-weighting based on species traits (Pillar et al. 2009). Given a set of *m* species spread across *n* communities, Beals (1984) smoothing predicts the probability of occurrence of every species *j* in each community *k*, estimated as the average of the pairwise co-occurrence probabilities of species *j* with those species actually present in community *k*. Fuzzy-weighting (Pillar et al. 2009) has some analogy to Beals smoothing but, instead of co-occurrence probabilities, it is based on trait similarities between species. Fuzzy-weighting results in a trait-based transformation of species composition in a metacommunity (Leibold et al. 2004) that can fully describe potential community composition regarding traits encompassing both convergence and divergence (Pillar et al. 2009). The correlation between these two matrices of predicted species composition should thus measure how well the traits used for fuzzy-weighting reflect the observed co-occurrence patterns. Hence, the objective of finding the set of functional traits mediating community assembly can be reduced to the task of developing an optimization algorithm that identifies the traits maximizing this correlation, together with an appropriate permutational test for significance.

To test our method, we generated data with known environmental filtering mechanisms and analysed how often our method correctly identified those traits involved in the simulated process of community assembly. Then, we applied the method to real plant community data, and checked whether it identified traits that can be considered relevant in driving species assembly in the studied communities.

## Methods

As input, the analysis uses community composition matrix **W** of sites by species, and matrix **B** of species described by traits. Here, we considered both simulated data generated under specified conditions and real data (see details in the following).

Beals smoothing (Fig. 1a) requires matrix **P** of pairwise probabilities of species co-occurrences, which is derived from the community composition matrix **W**:

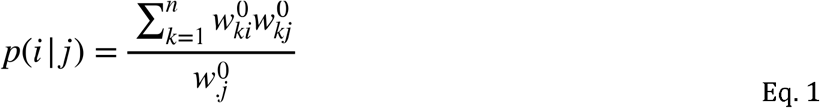

Where *p*_*i|j*_ is the probability of species *i* to occur in a community when species *j* is present, *w^0^*_*ki*_ and *w^0^*_*kj*_ are the incidences (0, 1) of species *i* and *j* in community *k*, and *w^0^*_*.j*_ is the total incidence of species *j* across the *n* communities in matrix **W**. Normalising W by its site-totals, to compute relative species abundances (**W**_p_), and multiplying it by **P** (Fig. 1a) results in Beals smoothed matrix **Y** of species by communities (Beals 1984; De Cáceres & Legendre 2008). In this definition, the target species was included for the estimation of their own probability of occurrence in a community (Beals 1984).

**Figure 1.**
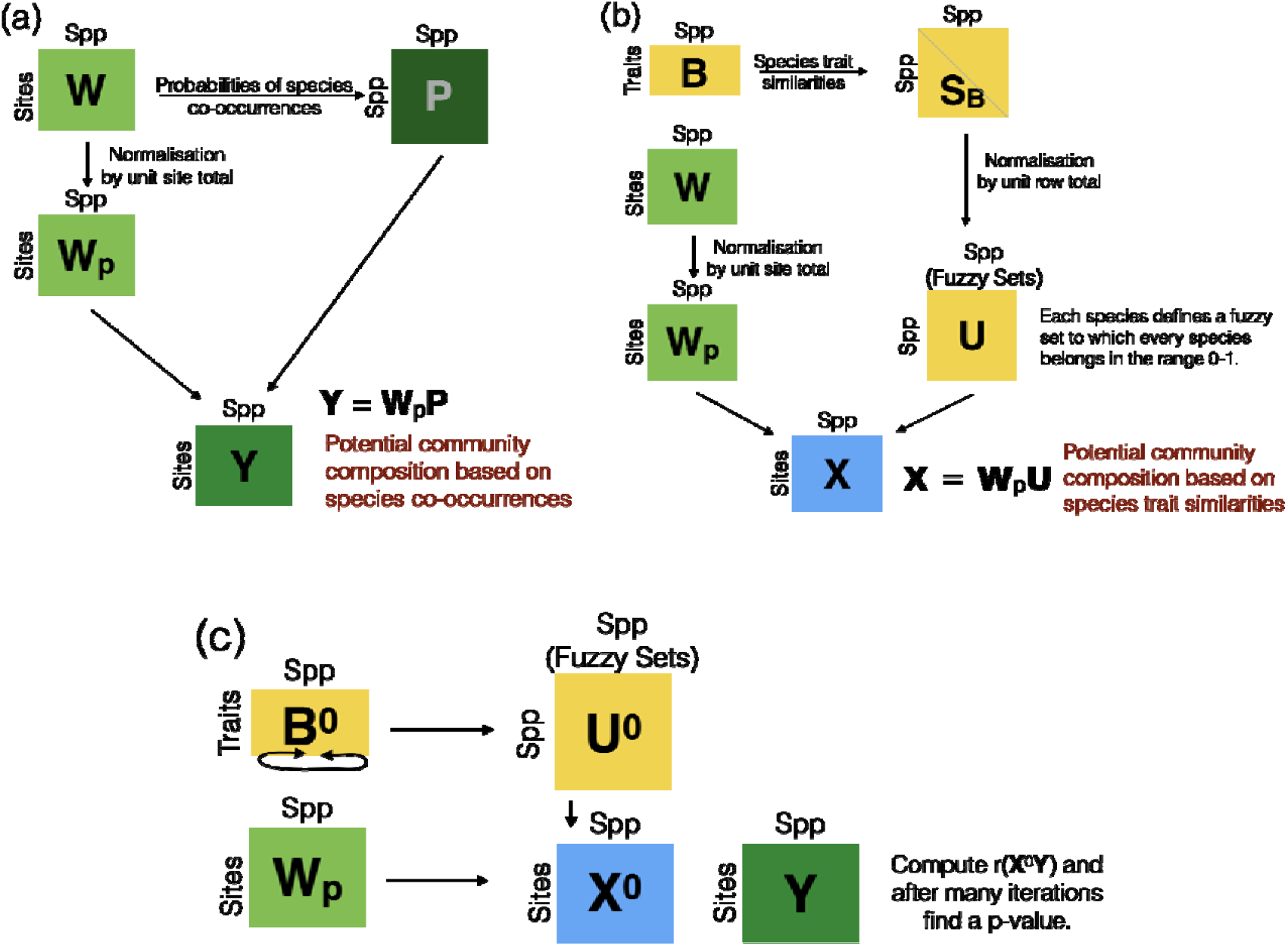
Data analysis steps for (a) Beals smoothing applied to the species composition matrix **W** to generate the matrix **Y**, (b) fuzzy-weighting applied to the species composition matrix **W** which, combined with the species traits in matrix **B**, generates the matrix **X**, and (c) permutation test for the significance of the matrix correlation r(**XY**) by permuting the columns of **B** (or **U**) generating B^0^ (or **U**^0^) and derived **X**^0^ = **W**_p_**U**^0^.

For the fuzzy-weighting of community composition in **W** (see Fig. 1b), the species probability of occurrence in a community is estimated based on the species’ trait similarities with other species observed in the same community (Pillar et al. 2009). For this task, considering the traits in **B**, a species by species similarity matrix **S** is computed by using the Gower similarity index (ranging 0-1). By normalising the rows of **S** by their row total, a matrix **U** is obtained whose elements define self-cross belongings between species (Duarte et al. 2016). Each column *j* of **U** defines a fuzzy set of species functionally similar to species *j*. The closer a given species is to species *j* in trait space, the higher is its degree of belonging to the fuzzy-set *j* and the better it can functionally represent the species *j*. Fuzzy-weighted community composition is computed by multiplying site-total standardised **W**_p_ by **U**, resulting in a communities by species matrix **X** (Fig. 1b). Each element in **X** is an estimation of the probability to find species *j* in community *k*, given the functional similarity of species *j* to the species actually occurring in community *k*.

To assess the correlation r(**XY**) between matrices **X** and **Y**, we used the Rd coefficient (Omelka & Hudecová 2013), which is a Pearson correlation coefficient of the Gower-centred pairwise distances (Gower 1966) based on **X** and **Y**, considering the full distance matrices. The closer Rd is to 1, the higher is the association between community distances in fuzzy-weighted species composition based on traits and those in potential composition based on species co-occurrences. The Rd correlation r(**XY**) can be interpreted as the degree to which the traits used in **X** reflect co-occurrence patterns in **Y**. We chose the Rd coefficient based on unsquared Euclidean distances because, compared to the Mantel correlation or the RV coefficient (Robert & Escoufier 1976), it can also detect non-linear relations between the matrices (Omelka & Hudecová 2013).

### Testing for significant traits

The significance of the Rd correlation r(**XY**) was tested under the null hypothesis that species assembly is unrelated to species traits (Pillar et al. 2009). This is achieved by keeping **W** and **Y** constant and permuting the columns of **B** (or, equivalently, of **U**) many times to allow the computation of a probability P(r(**X**^0^**Y**) ≥ r(**XY**)) (Fig. 1c). If the p-value is not larger than the a priori fixed error probability threshold α, r(**XY**) is deemed significant and we conclude that the trait or traits included in the definition of X has/have been relevant for community assembly. This permutation approach breaks all relations between the functional trait characteristics of the species and their presence or abundance in **W**, which has the following advantages: First, it controls for the fact that species composition (**W**) is used to derive the matrices at both sides of r(**XY**), thus it avoids bias that would result if permutations were done among sites in **X** or **Y**. Second, it avoids the source of bias described by Hawkins et al. (2017) affecting aggregated measures in community analysis; thus it conforms to the permutation solution described in Zelený (2018) for the analogous case of the community-weighted mean approach. Third, by keeping **W** and **Y** constant, any spatial or temporal autocorrelation in the compositional data will be incorporated in the null model, thus avoiding bias in the permutation testing (Pillar et al. 2009; Gotelli & Ulrich 2012).

This permutation procedure can be repeated by considering different subsets of traits for deriving fuzzy-weighted community composition in **X**. The trait or combination of traits maximizing r(**XY**), as long as its p-value is significant, is expected to be optimal for observational and experimental studies aiming to identify traits linked to hidden environmental factors in community assembly.

To select the optimal subset of traits, for the simulated data we considered the *p*-values generated according to Fig. 1c only, whereas for the real-world data we combined the permutation test with bootstrap resampling. Thus, since the real-world data are a sample, in addition to testing for significance, we calculated confidence intervals for the observed r(**XY**) for each trait or trait combination, and compared these across traits or trait combinations. For this, in each bootstrap iteration, the plots were resampled with replacement to obtain a bootstrap sample, which was then used to redefine **X**\* and **Y**\* with the selected plots and recalculate r(**X**\***Y**\*). We used the distribution of r(**X**\***Y**\*) across bootstrap samples to determine the 95% confidence interval of observed r(**XY**). Yet, as both **X** and **Y** are based on the same species composition **W**, they are expected to have non-zero r(**XY**) even if the trait combination used to build **X** plays no role in community assembly. Thus, we applied the permutational approach shown in Fig. 1c to compare r(**X**\***Y**\*) with a possible expected correlation r(**X**\*^0^**Y**\*) assuming the selected trait or traits has/have no role in community assembly. After a large number of bootstrap/permutation iterations, the probability P(r(**X**\*^0^Y) ≥ r(**X**\***Y**\*)) was the proportion of iterations in which r(**X**\*^0^**Y**\*) was larger than r(**X**\***Y**\*).

Finally, we used the 95% confidence intervals of each correlation r(**XY**) to compare and rank trait combinations. Ideally, we would examine iteratively every trait subset with 1 to *k* traits in **B** and the corresponding significance of the resulting r(**XY**). However, when the number of traits is large (e.g., >20), the number of possible combinations may become numerically unmanageable (e.g., 1,048,575 possible combinations for 20 traits). Therefore, we adopted a partial stepwise algorithm to efficiently explore the space of trait combinations and reduce computation demand, and we benchmarked the results with those of the analyses performed on simulated data with known assembly rules. The algorithm acts as follows: once computed r(**XY**) for each single trait, the traits resulting in significant r(**XY**) correlations were selected. We then repeated the procedure by considering all the pairwise combinations of traits being individually significant. If any pairwise combinations had an r(**XY**) significantly better than the best trait (i.e., whose 95% confidence intervals did not overlap with those of the best traits), we considered the pairwise combination having the highest and significantly better r(**XY**) as the new best. We then kept these two traits as fixed, while testing the effect of adding another trait, trying to find a new best. If no pairwise combination performed better than the best trait, we tested all possible three-way combinations, and checked if a new best could be found. We added one trait at the time until finding the optimal combination of traits. For each combination, we generated *p*-values using 999 random iterations of bootstrap/permutation plus one iteration for the observed r(**XY**).

### Analyses with simulated communities

To test whether our method is capable of discriminating relevant from non-relevant traits, we applied it to simulated plant community composition data. We generated data by modelling metacommunities (sets of plant communities) based on specified assembly mechanisms in which the underlying environmental factors were known. Then, we analysed the simulated data with the above-described method to identify the traits driven by these factors. This way we could check by means of type I error and power analyses whether the relevant traits for the assembled communities were correctly revealed.

We used a stochastic, individual-based model for simulating metacommunities stepwise from a pool of species and their functional traits (Pillar & Camiz 2020). At each step, the model predicts the arrival, establishment, and extinction of individuals belonging to each species, based on probability functions with specific parameters. We then analysed the metacommunity resulting after a given number of years (iterations). We generated different simulated metacommunities by specifying different combinations of trait numbers, environmental filtering parameters and species-level trait correlations (Appendix S1). The other parameters were set randomly. For each set of model parameters, we generated and analysed a total of 100 simulated metacommunities.

We explored three sets of simulation scenarios to assess whether the method can correctly identify the relevant traits in the simulated metacommunities, when confounding aspects related to correlation among traits and contrasting strengths and interactions between environmental filtering effects are in play. In the first case, we generated communities assuming two environmental factors and three functional traits. The first trait *t_1_* was directly dependent on *e*_*1*_, i.e., *e*_*1*_ -> *t*_*1*_, while *t*_*2*_ related directly to *e*_*2*_, i.e., *e*_*2*_ -> *t*_*2*_. An additional trait *t*_*n*_ was neutral with respect to the environment. We generated metacommunities under increasing magnitude of *e*_*1*_ -> *t*_*1*_, as given by the specified linear response parameters for environmental filtering, from 0 to 0.6, while fixing the effect of *e*_*2*_ -> *t*_*2*_ at 0.3. We used this basic scenario to explore both the effect of an interaction between environmental factors *e*_*1*_ and *e*_*2*_ on *t*_*1*_ (three levels: 0, 0.3, 0.5) and to explore the effect of the correlation between traits *t*_*1*_ and *t*_*2*_ (three levels, 0. 0.4, 0.8).

The second set of scenarios was similar to the first one, but we added a third trait *t*_*3*_ directly dependent on factor *e*_*1*_, i.e., *e*_*1*_ -> *t*_*3*_. In this case, both traits *t*_*1*_ and *t*_*3*_ were affected by the same factor *e*_*1*_, but while the strength of the effect *e*_*1*_ -> *t*_*1*_ varied from 0 to 0.6, the effect *e*_*1*_ -> *t*_*3*_ was fixed at 0.3. As in the first set of scenarios, we also examined the effect of an interaction between factor *e*_*1*_ and *e*_*2*_ on *t*_*1*_, and of pairwise correlations between traits *t*_*1*_, *t*_*2*_ and *t*_*3*_.

In the third set of scenarios, we varied the effect *e*_*1*_->*t*_*1*_ from 0 to 0.6, as above, but progressively included also the effect of additional environmental factors on respective functional traits (i.e., *e*_*2*_->*t*_*2*_; *e*_*3*_->*t*_*3*_; *e*_*4*_->*t*_*4*_), all with a magnitude of 0.3. In all simulations, a neutral trait *t*_*n*_ was added with the purpose of testing type I error. Factor interaction effects and pairwise trait correlations were set to zero in these scenarios.

The analysis allowed evaluating the power of the method, i.e., the proportion of metacommunities in which traits involved in the simulated assembly mechanisms were correctly identified as being significant, i.e., when the test with the simulated metacommunity resulted in P(r(**X**^0^**Y**) ≥ r(**XY**)) ≤ 0.05. It also allowed evaluating type I error or the accuracy of the method, i.e., the proportion of metacommunities in which neutral traits (*t*_*n*_ and also when the effect *e*_*1*_ -> *t*_*1*_ was set to zero) were incorrectly identified as relevant. For the simulated data, significance was evaluated for traits considered individually and for all possible trait combinations.

### Analyses with real communities

To test whether our method is helpful in highlighting relevant traits in a real-world dataset, we used data on dry calcareous grasslands vegetation in Germany. Such grasslands belong to the *Festuco-Brometea* class (Mucina et al. 2016) and are coded “E1.2a Semi-dry perennial calcareous grassland” in the European Red List of Habitats (Dengler et al. 2017). The dataset was previously used in a continental survey (Willner et al. 2019). Here we analysed a subsample of 565 plots randomly taken from the original data (see map in Appendix S2), and including 488 species. We combined compositional data (square-root transformed percentage cover) with species trait information for 49 traits (Appendix S3) from the BIOLFLOR (Klotz et al. 2002) and TRY databases (Kattge et al. 2011; Kattge et al. 2020). The TRY data, which included 16 traits, were gap-filled, as described in (Shan et al. 2012; Fazayeli et al. 2014; Schrodt et al. 2015; Bruelheide et al. 2019). Trait coverage was complete except for pollination, leaf persistence, sclerophylly, and succulence, for which the species with functional trait information accounted for an average of at least 96.5% of the plot total cover across the plots in our sample (Appendix S3).

The r(**XY**) correlations were calculated for all traits, first trait by trait, and then testing the traits with highest r(**XY**) in combination based on the stepwise algorithm described above. This allowed to identify the optimal trait subset, i.e., the combination of traits with the maximum relevance for the assembly of these grassland communities. We used principal components analysis (PCA) based on pairwise trait correlations to identify the main trends of trait variation at the species level.

To illustrate how well the selected traits reflected community composition, we calculated a PCA of the dry grassland data based on the covariance of fuzzy-weighted composition (**X** matrix). The principal components were then passively projected on another PCA based on the covariance of Beals’ smoothed composition (**Y** matrix). Also, the CWMs of all relevant traits were projected on this ordination space based on their Pearson correlations with the principal components. In addition, to explore environmental explanations for the observed community trait composition, we compiled annual mean temperature and annual mean precipitation from CHELSA, V1.1(Karger et al. 2017) and assigned these values to the plots with a 30 arcsec resolution. Also, two soil variables (soil pH and content of soil organic carbon) were extracted from the SOILGRIDS project (https://soilgrids.org/, licensed by ISRIC—World Soil Information), downloaded at 250 m resolution and then resampled using the 30 arcsec grid of CHELSA. These environmental data were also projected on the ordination space based on their Pearson correlations with the principal components of the community composition.

## Results

### Simulated communities

In the first set of scenarios (Fig. 2, top, leftmost panel), the proportion of simulated metacommunities with a significant r(**XY**) correlation taking trait *t*_*1*_ alone expectedly increased when the factor effect *e_1_* on trait *t_1_* increased beyond zero, and reached 100% power with the strongest effect. However, as the effect of factor *e*_*1*_ on trait *t*_*1*_ increased, the power to detect a significant r(**XY**) for trait *t*_*2*_ alone was suppressed. In addition, the method correctly indicated that the proportion of simulated metacommunities with significant r(**XY**) for *t*_*n*_ alone was low and close to the nominal α threshold of 0.05, i.e. the type I error was not inflated. However, considering combinations of traits showed that all two-trait combinations involving the neutral trait *t*_*n*_ returned significant r(**XY**) at similar power to the one obtained when considering traits *t*_*1*_ or *t*_*2*_ alone. This is clearly misleading considering that tn was not under environmental filtering in community assembly. We took this result as evidence for the need to only test combinations of traits which produced a significant r(**XY**) when taken individually.

**Figure 2.**
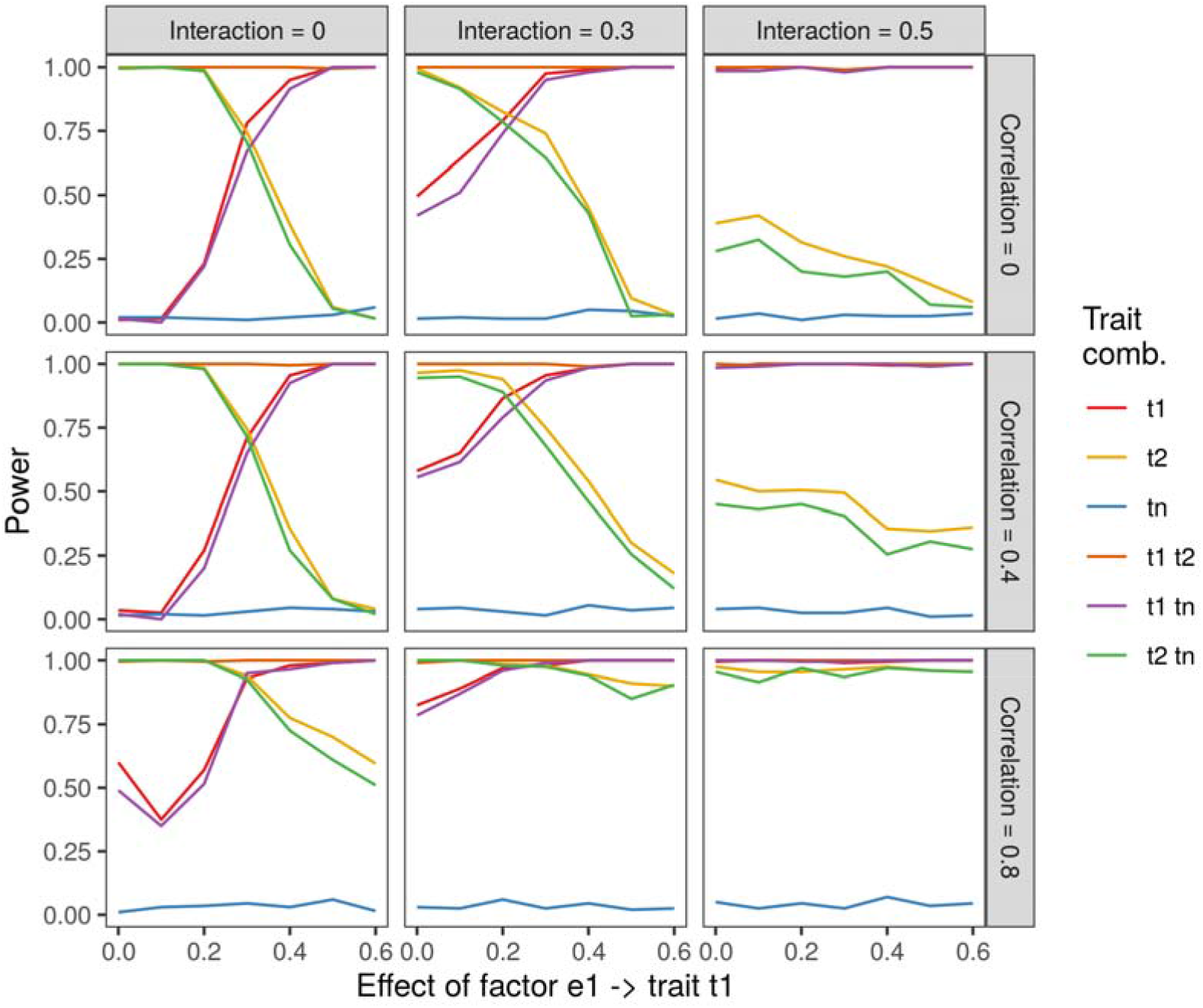
Simulated-data power profiles of Rd matrix correlation r(**XY**) between community distances based on trait-based fuzzy-weighted (**X**) and Beals-smoothed (**Y**) species composition for metacommunities with increasing strength of factor effect *e*_*1*_ on trait t1, and varying the magnitude of the *e*_*1*_ x *e*_*2*_ interaction, and the strength of the pair-wise correlations between traits *t*_*1*_ and *t*_*2*_ (Scenario 1). Power (vertical axis) is the proportion of simulated metacommunities for which the P-value for r(**XY**) found by permutation was not larger than a threshold of 0.05. The graphs show traits considered individually and different trait combinations defining fuzzy-weighted species composition. Further details on the set parameters for community assembly simulations are in Appendix S1.

Furthermore, as the effect of factor interaction *e*_*1*_ x *e*_*2*_ on trait *t*_*1*_ increased (Fig. 2, top panels), the relevance of *t*_*1*_ was high irrespective of how low the factor effect *e*_*1*_ was on the same trait. The power to detect a significant r(**XY**) for *t_2_* alone was even more strongly suppressed with increasing interaction *e*_*1*_ x *e*_*2*_ on trait *t*_1_ (Fig. 2, mid and right column of panels). However, when the correlation between traits *t_1_* and *t_2_* increased (Fig. 2, mid and bottom panels), the suppression of trait *t*_*2*_ by trait *t*_*1*_ was not any longer evident.

The suppression effect between traits can be better examined in the second set of scenarios (see results in Appendix S4). Similarly, to what shown in Fig. 2, in the absence of factor interaction and trait correlation the detection of trait *t*_*2*_ as relevant in community assembly was progressively suppressed by *t_1_* when the filtering effect of factor *e*_*1*_ increased. However, trait *t*_*3*_, which in this scenario is filtered by the same factor e1, was much less suppressed as the filtering effect on trait *t_1_* increased, i.e., became more limiting for the establishment and the survival of plant individuals. Yet, under increasing strength of the interaction *e*_*1*_ x *e*_*2*_ on trait *t*_*1*_, the power to detect a significant r(**XY**) for *t_3_* alone decreased. Further, similar to the first set of scenarios, increased pairwise correlation at the species level between traits *t*_*1*_, *t*_*2*_ and *t*_*3*_ reduced such a suppression effect. As before, the type I error was not inflated regarding the neutral trait *t*_*n*_ taken alone.

In the third set of scenarios, we analysed whether the performance of our method is influenced by the number of traits involved in community assembly (Fig. 3). The simulations based on three traits generated power graphs with a similar pattern compared to the ones based on four or five traits. In all cases, trait *t_1_* was filtered under increasing factor effect *e*_*1*_, trait *t*_*n*_ was always neutral and the other traits were under a fixed, intermediate factor effect. As in Fig. 2, the analysis of the r(**XY**) using trait combinations including the neutral trait tn would be as relevant as using the other non-neutral traits alone.

**Figure 3.**
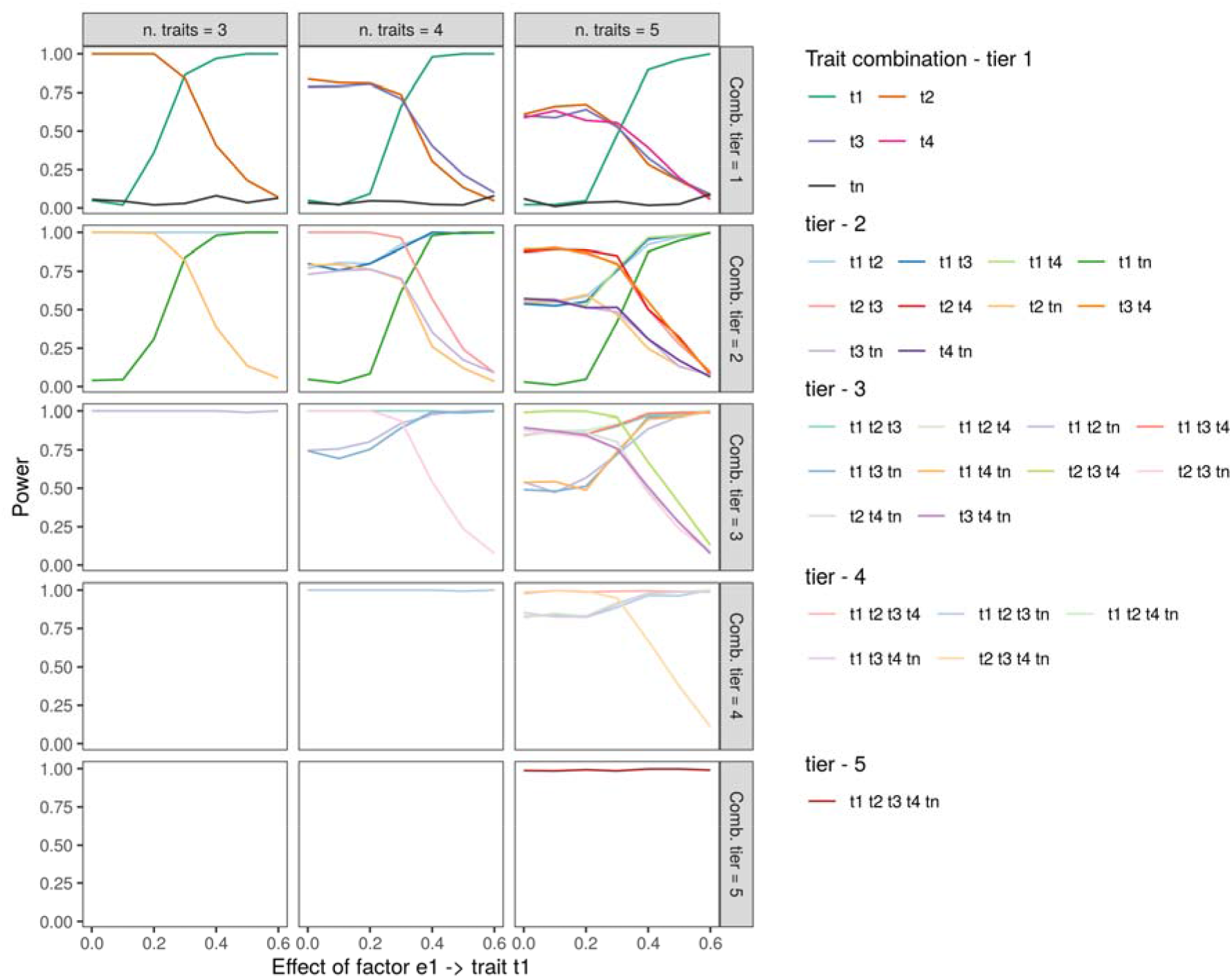
Simulated-data power profiles of Rd matrix correlation r(**XY**) between community distances based on trait-based fuzzy-weighted (**X**) and Beals-smoothed (**Y**) species composition with increasing strength of factor effect *e*_*1*_ on trait *t*_*1*_, and increasing the number of traits used in simulating metacommunities (Scenario 3). Power (vertical axis) is the proportion of simulated metacommunities for which the p-value found by permutation was not larger than a threshold of 0.05. The number of traits ranged from 3 to 5 (left to right panels), with one trait *t*_*n*_ always being neutral. Traits are either shown individually (top row), or in combinations (from two to five, top to bottom rows) to improve visualization.

### Real communities

When applying the approach to German dry calcareous grasslands, seven out of the 49 traits returned a significant r(**XY**) when taken one by one: sclerophylly, plant height, specific leaf area (SLA), nanophanaerophyte and hemiphanaerophyte growth-forms, flowering duration, and vegetative propagation through fragmentation. Taken singularly, sclerophylly was the trait that best explained community assembly (Fig. 4).

**Figure 4.**
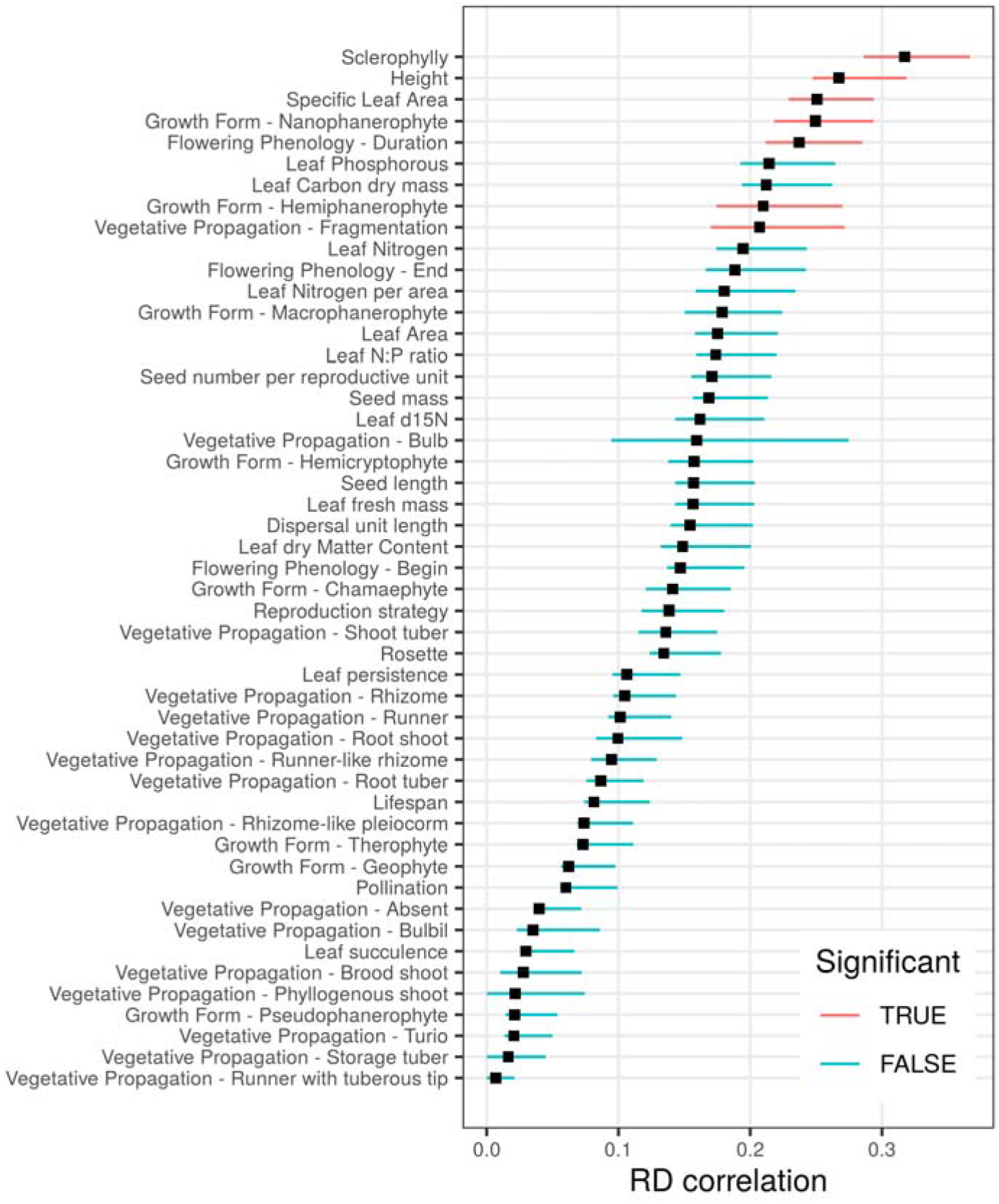
Rd matrix correlation r(**XY**) between community distances based on trait-based fuzzy-weighted (**X**) and Beals-smoothed (**Y**) species composition, when considering one trait at the time. The observed r(**XY**) was deemed significant (at p-value ≤ 0.05, one-sided) when it was greater than the respective correlation coefficient calculated using permuted species traits in at least 97.5% of the bootstrap samples. The segments represent 95% bootstrap confidence intervals of the observed r(**XY**); in red are the traits with significant r(**XY**), in blue are the non-significant ones.

Increasing iteratively the number of traits used to calculate the **X** matrix, resulted in a progressive increase in r(**XY**), although the confidence intervals of the regression coefficients were mostly overlapping (Fig. 5). When considering pairwise combination of traits, the combination sclerophylly and flowering duration, returned a significantly higher r(**XY**), compared to sclerophylly alone. There were no three- and four-way combinations of traits significantly improving the r(**XY**) compared to the sclerophylly-flowering duration couple (for details see Appendix S5). Only when considering five traits together, the improvement in r(**XY**) became significant: beside sclerophylly and flowering duration, the other traits composing this combination of traits were plant height, SLA, and propagation by fragmentation. We defined this as being the optimal combination of traits for predicting fuzzy-weighted species composition related to species co-occurrences, as no additional increase in dimensionality resulted in a significant improvement in r(**XY**) (Fig. 5).

**Figure 5.**
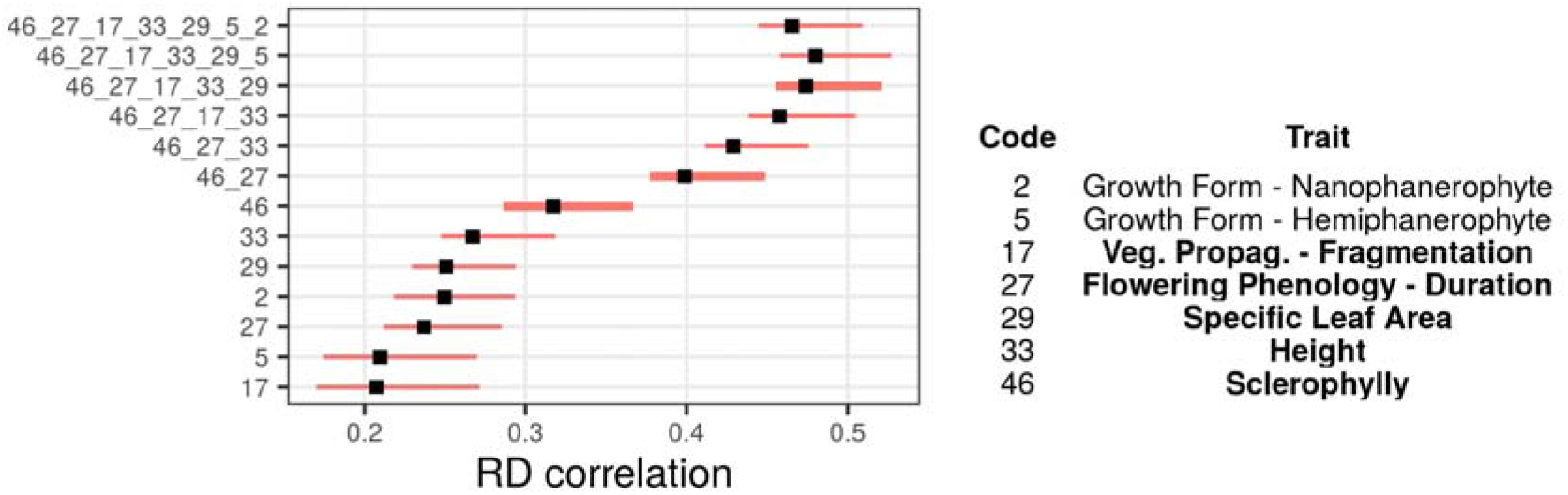
Rd matrix correlation r(**XY**) (black squares) and confidence interval (red lines) between community distances based on trait-based fuzzy-weighted (**X**) and Beals-smoothed (**Y**) species composition, when progressing in tiers (bottom to top) based on a selected subset of traits. Only the seven significant traits defining fuzzy-weighting alone (see Fig. 4) were considered. At each tier, we tested the effect of adding a new trait to the best combination of the previous tier, and only show the best result. We used thick lines for traits or trait combinations providing a significant (p<0.05) improvement with respect to the best solution at the previous tier(s). Detailed results are shown in Appendix S5.

The analysis of the trait correlations at the species level (Appendices S6-S7) revealed two main axes of independent trait variation, one reflecting the leaf economics spectrum (SLA vs. sclerophylly), which was also associated with hemiphanerophyte growth form and propagation by fragmentation, and the other the size spectrum (plant height), which was also associated with nanophanerophyte growth form and flowering duration. However, uncorrelated traits at the species level were not necessarily also uncorrelated at the community level. For example, while at the species level plant height was uncorrelated to sclerophylly (r = −0.06) and fragmented vegetative propagation (r = −0.07), their corresponding CWM values showed considerable Pearson correlations (−0.44 and 0.31, respectively Appendix S8).

The five traits that were identified as the most relevant ones (Fig. 5), and the so defined principal components of fuzzy-weighted composition (FW-PCs, Appendix S9) reflected different dimensions (PCs) of Beals smoothed community composition, as shown in Fig. 6 (see correlations in Appendix S10). FW-PC2 reflected an increasing representation of the nanophanerophyte growth form vs. decreasing flowering duration and was mostly correlated to the first principal component (PC1) of the Beals smoothed community composition (27.7% of total variation). FW-PC1 reflected the leaf economics spectrum (SLA vs. sclerophylly) and was correlated also to PC1 but mostly to PC3 of the Beals smoothed community composition (11.2% of total variation). FW-PC3 was only (weakly) correlated to PC4 but did not reflect any trait in particular. Yet, the links between the FW-PCs, the traits and the PCs of the Beals smoothed community composition become clearer by examining the two-dimensional ordination spaces. In the space defined by PC1 and PC2, two diagonal axes can be identified, one reflecting FW-PC1 and the other FW-PC2, both representing different traits. The size spectrum (height) was captured by both FW-PC1 and FW-PC2. Finally, the available potential environmental predictors presented weak correlations with the first four principal components, being highest for mean annual precipitation (−0.386 with PC1, Fig. 6, Appendix S10).

**Figure 6.**
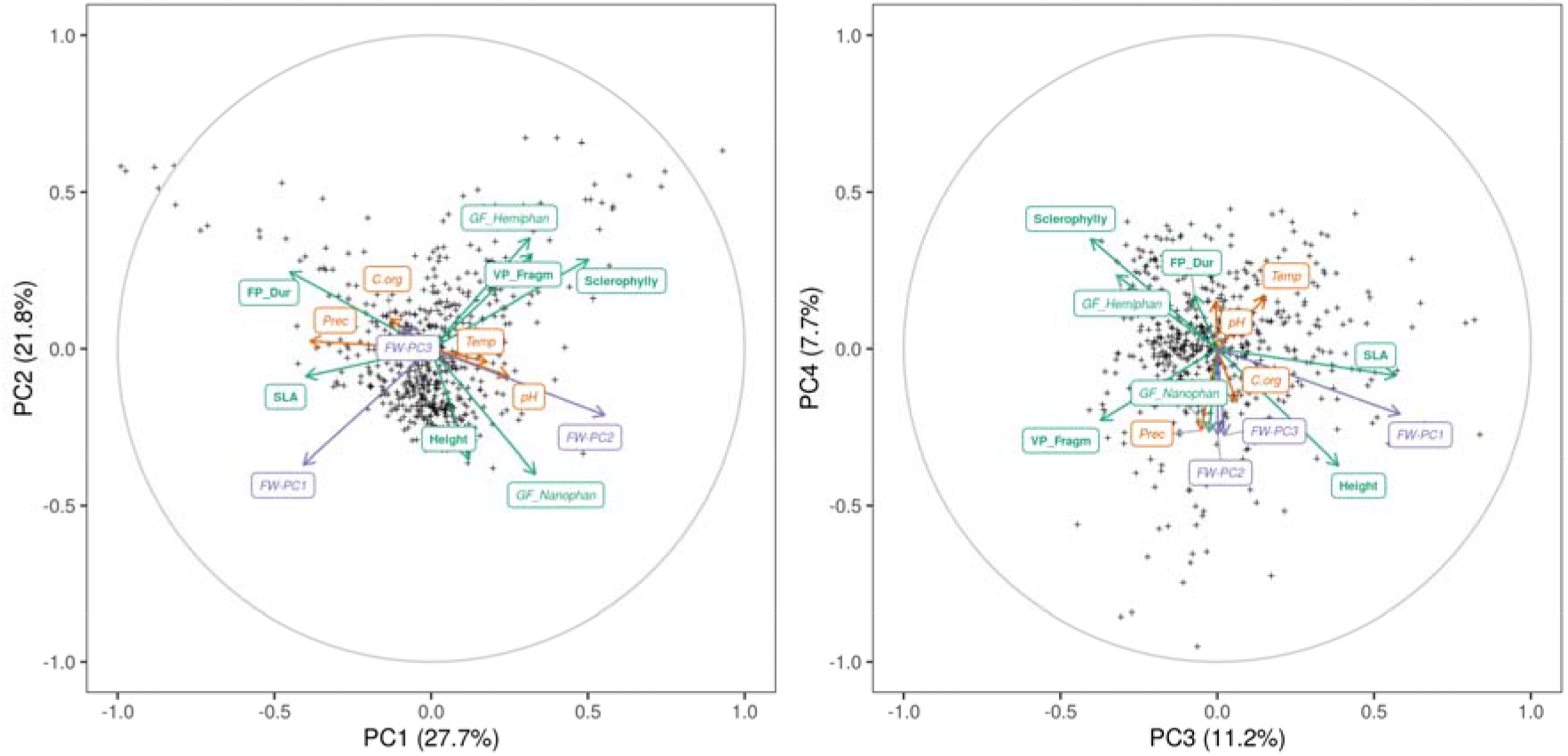
Principal component analysis of German dry grassland plots based on the species variance-covariance matrix of Beals’ smoothed composition (**Y** matrix), shown as cross symbols. The CWMs for the traits with a significant Rd matrix correlation r(**XY**) in Fig. 4, in green, the principal components based on the fuzzy-weighted composition defined by these traits (FW-PC1, -PC2, -PC3, Appendix S9), in purple, and four environmental variables, in orange, are projected on the ordination space according to their Pearson correlations with the PCA axes (see correlations in Appendix S10). The five traits identified in Fig. 5 as the best combination of traits are shown in bold green fonts. See Appendix S11 for the scatterplots with the species.

## Discussion

How to identify those functional traits driving community assembly when relevant environmental factors are unknown? Answering this question is crucial to improve our predictions on how ecological assemblages will change in the face of global change (Newbold 2018). Here, we developed a method to identify the functional traits mediating community assembly, which does not rely on measuring the actual the environmental gradients ultimately driving it. Our approach relies on the comparison of two alternative ways of predicting how species are likely to occur in a given community: Beals’ smoothing of species co-occurrences probability (Beals 1984) and fuzzy-weighting of functional traits (Pillar et al. 2009). The method comes with an optimization algorithm able to efficiently explore the trait combination space, and derives unbiased significance values and confidence intervals using permutation.

The results with the simulated data show that our method proved capable of identifying the most relevant trait combinations mediating the assembly of biological communities along gradients. The power of our analysis quickly increased to 100%, when the magnitude of the main environmental filtering effect, specified as a linear parameter relating the factor to the expected trait values at the community level in the metacommunity model that generated the data, was greater than 0.3. This suggests that the method might be sensitive enough to detect the most important traits related to discriminant environmental factors in real-world situations. Furthermore, our approach proved sufficiently robust against the inclusion of non-relevant traits, being the type I error always close to the nominal levels, as well as against confounding factors related to interactions between environmental gradients, and correlation among traits.

The results with the simulated data, however, also indicated that only those traits found relevant when taken individually should be retained in the analysis and tested in combination with other equally relevant traits. In other words, considering correlation and *p*-values *per se*, was not sufficient to discriminate trait combinations which include irrelevant traits. For the real data, we solved this problem by using bootstrap to calculate the confidence intervals of our matrix correlation coefficients, and by adopting a partial stepwise algorithm only considering combinations of traits that were relevant when taken individually. This way, we could reliably ascertain that a combination of, e.g., two traits, was significantly better than any of the two traits taken singularly. And yet, our optimization algorithm remained sufficiently flexible to be adapted to situations in which the examination of every combination of relevant traits would be unfeasible.

Our results using simulated metacommunity data demonstrated a suppression effect among traits in their role in community assembly, suggesting that traits under stronger filtering effects tend to mask traits that are weakly filtered. Suppression may arise from the obvious fact that the units being filtered are not traits but whole organisms, whose traits cannot be physically disentangled according to trait responses to different factors. Under such filtering effect, the most limiting trait (Sih & Gleeson 1995; Gorban et al. 2011) likely suppresses less limiting traits. However, suppression is stronger between traits that are filtered by different, independent environmental factors than between traits that are filtered by the same factor. Correlation among traits, on the other hand, reduces such suppression effects.

Models are useful but offer simplified representations of real systems. Thus, models should always be confronted with or complemented by the analysis of real data (Noy-Meir & van der Maarel 1987). We believe this approach was successful here. While there is no way to disentangle all the environmental factors that drive the community composition of the whole range of dry calcareous grasslands in our study system, the identification of the five most relevant traits allows some conclusions on the underlying processes. Three of the five traits are part of the two main spectra of global plant forms and functions at the species level (Díaz et al. 2016). While plant height reflect the size spectrum, SLA and sclerophylly represent the leaf economics spectrum (Wright et al. (2004).

Plant height, on the one hand, points to succession as a key factor in community assembly of dry grasslands. Indeed, abandonment of grazing and mowing favours tall grasses, shrubs and trees, i.e. plants of higher stature. Taller species indicate ongoing secondary succession, which is a major threat for dry grasslands (Kahmen & Poschlod 2004; Burrascano et al. 2016). We found that the successional gradient is reflected by the first and second dimensions of the fuzzy-weighted composition based on the five key traits, which supports the result from experiments that revealed land use intensity and time since abandonment as main drivers of trait composition of dry grasslands (Moog et al. 2002). On the other hand, the leaf economics spectrum, characterized by specific leaf area (SLA) versus sclerophylly (Wright et al. 2004), forms a second gradient, yet not completely independent of the successional one. In our communities, the ability to propagate through fragmentation coincides with the leaf economics spectrum gradient because this trait is represented in slow-growing perennial species that fragment with age. In dry grasslands, the leaf economics spectrum reflects the gradient in both nutrient and water supply, along which different communities, alliances and orders are distinguished (Royer 1991; Jandt 1999; Willner et al. 2019). However, the overall nutrient availability, especially of N and P supply in these grasslands is low, making them rather stressful habitats, home to many specialist species adapted to these specific conditions (Gilbert et al. 2009; Ceulemans et al. 2011). These conditions also favour the hemiphanerophytic life form, (i.e. resting buds are situated on woody shoots).

These explanations might give the impression that the five key traits follow clear environmental gradients of easily measurable variables, yet the real-world situation is much more complex. While to some degree the plant height and leaf economics spectra follow macroclimatic gradients and result in different species pools of dry grasslands (see the map of the species pools in Bruelheide et al. (2020)), microclimate might strongly deviate from macroclimate (Bruelheide & Jandt 2007; Burrascano et al. 2013). Similarly, topographical conditions and soil depth have strong impacts on water availability, resulting in small-scale variation of communities (Leuschner 1989). This is illustrated by one of our five traits of the optimal combination, that is flowering period duration. The CWM of this trait was correlated with neither the community trends related to height nor to the leaf economics spectrum. This is consistent with the results reported by Bouchet et al. (2017): while flowering period duration showed a strong relationship to community trait composition, was not related to successional age. We would assume that flower duration indicates a combination of environmental factors that are usually hidden behind the main effects of these factors. Flower production depends on availability of resources and is supported by warm and wet conditions (Craine et al. 2012). These conditions occur in early successional stages with an open vegetation structure where deeper soils provide an above-average resource supply. In trait space, these particular micro-environmental conditions would promote a combination of low-stature growth close to the ground (small height) with acquisitive leaf traits (high SLA), to both of which flower period duration was moderately related.

While microclimate and soil depth are measurable, other additional factors adding to the complexity of dry grassland community assembly are not. In particular, historical factors are hidden in the present-day community assembly. For example, traditional shepherding between the 15^th^ and 20^th^ century has strongly affected species composition of calcareous grasslands (Poschlod & WallisDeVries 2002). There might be further hidden factors driving community trait composition, about which we can only speculate. For example, resource supply of dry calcareous grasslands can vary at very fine scales (Regan et al. 2014). This is both caused by a large variation of microsite soil conditions at small distances but also by heterogeneous effects of grazing. Overall, it becomes apparent that in real-world situations community composition is not driven by a single trait-environment relation, but a complex of different traits that are only partly related to known environmental factors.

Although trait divergence patterns may also arise in community assembly (Mason & Wilson 2006; Wilson 2007; Pillar et al. 2009), we did not examine the ability of our method to faithfully reveal relevant traits linked to biotic and/or abiotic factors causing trait divergence in the simulated community assembly. Yet, as the fuzzy-weighting adopted in our method integrates trait similarities at the species level fully into species composition matrix **X** at the community level (Pillar et al. 2009), we expected that relevant traits would be revealed irrespective of the actual mechanism, whether it generated trait convergence, trait divergence or both.

The method we proposed here successfully identified the relevant traits mediating community assembly, without relying on the measurement of the environmental factors responsible for the restrictions imposed on the species co-occurrence patterns. Trait-environment relations affecting community assembly (Keddy 1992; Wilson et al. 1999; Götzenberger et al. 2012) leave persisting marks in the patterns of species co-occurrences. These marks are revealed by our approach. Considering that individuals within species tend to be more similar to each other than between species (Kazakou et al. 2014; Siefert et al. 2015), by relating species traits to species co-occurrence in communities, our method is able to identify the traits most likely affected by those trait-environment relations, even when the environmental factors are hidden, unknown, or not easily measurable. Going beyond the reliance on measured environmental factors, our method is particularly promising in those domains where obtaining a set of consistent and comprehensive environmental measurements is unfeasible. We think specifically to analyse large biodiversity databases of co-occurrence data (Bruelheide et al. 2018; Bruelheide et al. 2019), where the use of our method might be instrumental to reveal the key traits underlying the geographical distribution of ecological communities, so to better infer the key ecological gradients behind these patterns.

## Data availability statement

The metacommunity simulation model and the power and accuracy analyses are implemented in the package SYNCSA, available at http://ecoqua.ecologia.ufrgs.br/SYNCSA.html. The R script used for the analysis of the grassland data is available at https://git.idiv.de/sPlot/hidden. The dry grassland vegetation plot data was extracted from the GVRD database available at https://www.givd.info/ID/EU-DE-014, and the sampled relevés are the ones listed in Appendix S12.

## Acknowledgements

This paper was mostly developed during a research visit of V.P. to the German Centre for Integrative Biodiversity Research (iDiv) Halle-Jena-Leipzig. In our analyses we used the iDiv High-Performance Computing (HPC) cluster, for which we in particular acknowledge the support of Christian Krause. The species trait data provided by the BIOLFLOR and TRY databases are acknowledged.

## Author contributions

V.P. conceived the method, with S.C. and H.B. contributions; V.P. and S.C. devised and implemented the metacommunity simulation model; V.P. and F.M.S. implemented computation tools and performed the analyses. U.J. curated the dry grassland data. All authors discussed the results and contributed on the manuscript.

## List of Appendices

Appendix S1. Parameters set for the simulation of metacommunities used for the analyses shown in Fig. 2.

Appendix S2. Location of the grassland plots from the dataset of Willner et al. 2019 stored in the GVRD database.

Appendix S3. Traits used for the analysis of dry calcareous grassland communities.

Appendix S4. Simulated-data power profiles of matrix correlation r(**XY**) between community distances based on trait-based fuzzy-weighted and Beals-smoothed species composition for metacommunities.

Appendix S5. Numerical results for best solutions shown in Figure 5.

Appendix S6. Principal components analysis (PCA) of species on the basis of the matrix of species by the traits found relevant in community assembly (Fig. 5).

Appendix S7. Pearson correlation matrix between traits at species level.

Appendix S8. Pearson correlation matrix between CWMs at community level.

Appendix S9. Principal components analysis (PCA) of dry grassland plots based on the fuzzy-weighted composition (**X** matrix).

Appendix S10. Pearson correlation coefficients between principal components of Beals’ smoothed composition and fuzzy-weighted composition, CWMs and environmental variables.

Appendix S11. Principal component analysis of dry grassland plots based on Beals’ smoothed composition (Y matrix), showing the species.

Appendix S12. Identification numbers of the relevés extracted from the GVRD database.

Appendix S13. Species found in dry calcareous grassland relevés extracted from the GVRD database.

