## Appendices for "Revealing the functional traits that are linked to hidden environmental factors in community assembly"

Supporting information to the paper Pillar, V.D. et al. Revealing the functional traits that are linked to hidden factors in community assembly, *Journal of Vegetation Science.*

**Appendix S1**. Parameters set for the simulation of metacommunities used for the analyses shown in Fig. 2. In (a) the parameters specify trait correlations and the levels of environmental filtering, i.e., community responses to ecosystem conditions that were fixed with different values within the indicated ranges for type I and power evaluations of the correlation r(**XY**). In (b) the parameters varied randomly within the specified ranges and were not relevant for type I and power evaluations. In (c) explanations are provided (further details will be found in Pillar & Camiz 2020).

(a) Parameters set within the indicated ranges for evaluating type I error and power of r(**XY**):

Specified trait correlation matrix (3 traits x 3 traits). Trait data generated with uniform distribution

| | $t_1$ | $t_2$ | $t_n$ |
|---|---|---|---|
| $t_1$ | 1 | 0 to 0.8 | 0 |
| $t_2$ | 0 to 0.8 | 1 | 0 |
| $t_n$ | 0 | 0 | 1 |

Parameters for linear models of community responses to ecosystem conditions (3 traits by 2 ecosystem predictor variables, and their interaction):

| | $e_1$ | $e_2$ | $e_1 \times e_2$ |
|---|---|---|---|
| $t_1$ | 0 to 0.6 | 0 | 0 to 0.6 |
| $t_2$ | 0 | -0.3 | 0 |
| $t_n$ | 0 | 0 | 0 |

(b) Parameters that were either fixed or varied randomly within the indicated ranges for every simulated metacommunity irrespective of the settings in (a):

| *Parameters set per metacommunity:* | *Fixed* | *Min.* | *Max.* |
|---|---|---|---|
| Number of species in the species pool | 150 | | |
| Number of communities in the metacommunity | 200 | | |
| Correlation level between environmental factors (2 factors x 2 factors) for generating environmental data (uniform distribution) | 0 | | |
| Maximum average number of individuals per community | 200 | | |
| Number of iterations (cycles of colonization and extinction) per year | 50 | | |
| Duration of the simulation | 100 | | |
| Site geographical coordinates | | 0 | 100 |
| *Parameters set per species and metacommunity:* | | *Min.* | *Max.* |
| Centre of antisymmetry of the logistic function for the dispersal kernel | | 0.1 | 0.9 |
| Negative slope of the logistic function for the dispersal kernel | | -5 | -2 |
| Centre of antisymmetry of the logistic function for survival probability | | 0.1 | 0.4 |
| Negative slope of the logistic function for survival probability | | -5 | -2 |
| Centre of antisymmetry of the logistic function for age-related survivorship curves | | 0.1 | 0.9 |
| Negative slope of the logistic function for age-related survivorship curve | | -5 | -2 |
| Centre of antisymmetry of the logistic function for density dependence | | 0.6 | 0.9 |
| Negative slope of the logistic function for density dependence | | -5 | -2 |
| Lifespan of individuals | | 1 | 50 |
| *Parameters set per species, trait and metacommunity:* | | *Min.* | *Max.* |

| | | | |
|---|---|---|---|
| Centre of antisymmetry of the logistic function for survival probability | | 0.1 | 0.4 |
| Negative slope of the logistic function for survival probability | | -5 | -2 |

(c) *Modus operandi*

For each simulated metacommunity, a matrix of species by traits with the specified correlation structure in (a) and a matrix of community sites by random uncorrelated environmental factors were generated. For the aims of our simulations, the locations of the community sites were set at random and independently from the environmental factors. Species colonization and extinction events were defined according to the probabilities of individuals arriving, establishing and dying at each community site. These probabilities were all defined by a logistic function with centre and slope parameters set randomly for each simulated species within specified ranges. That is, for seed arrival, the probabilities for each species were a function of the spatial distance, standardized to unit maximum, of the community site to the nearest seed source in the metacommunity. For environmental-driven survival, the probabilities were a function of the similarity between the individual and the expected community-weighted mean trait for each trait, and this in turn was a function of the environmental factor values at the site based on specified linear response parameters for environmental filtering. For density-dependent survival and mortality, the probabilities were a function of the total abundance of individuals relative to the maximum set number of individuals in the community, and for age-related mortality, the probabilities were a function of the individual age relative to the species expected longevity, which was set at random within the specified range. After a large number of years (100) with the specified number of 50 cycles of individual colonization and death per year, the simulated metacommunity data was analysed and the functional variation patterns compared to the expected ones.

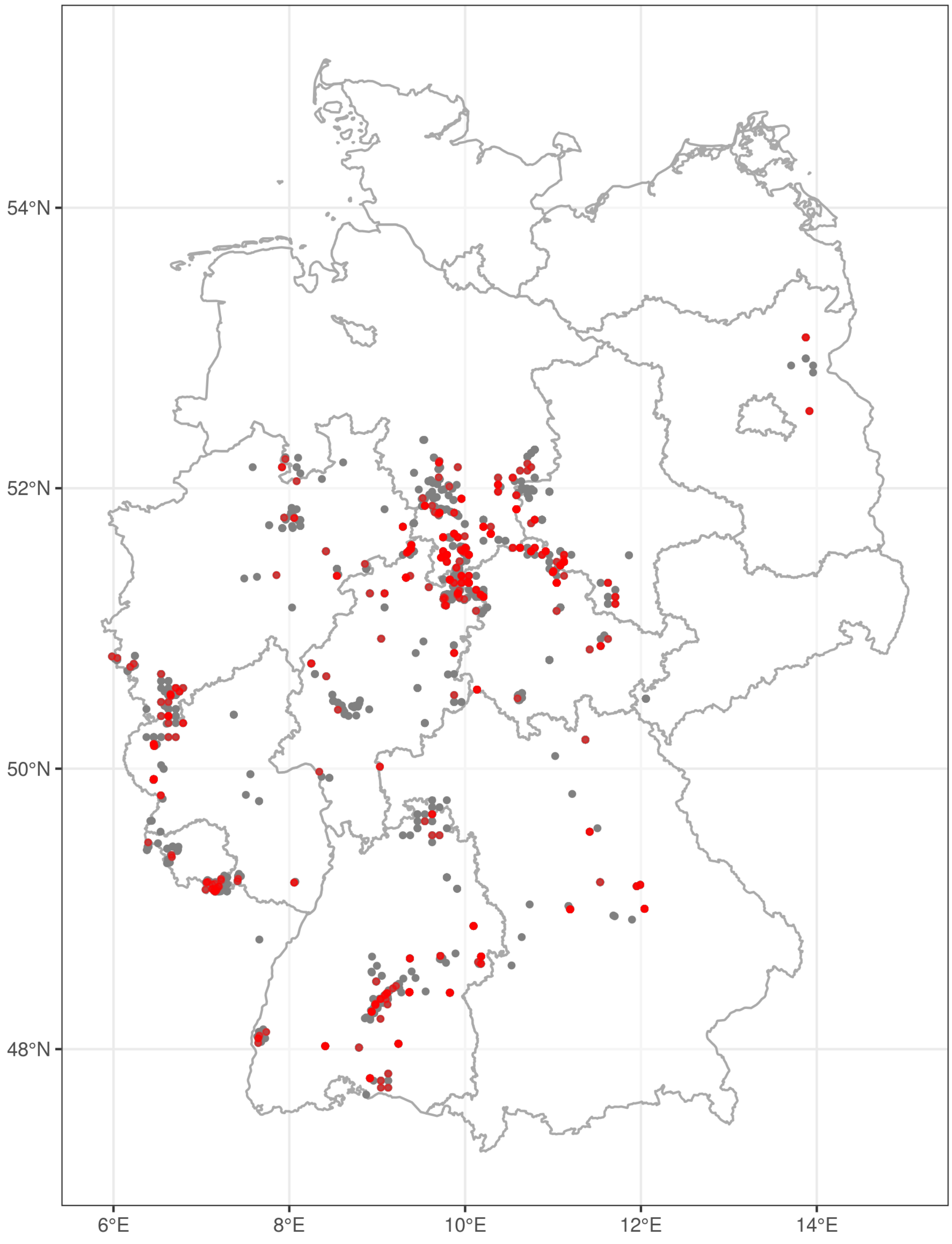


**Appendix S2**. Location of the grassland plots from the dataset stored in the GVRD database, which were used for the European classification of dry grasslands (Willner et al. 2019). The 565 selected plots for our analysis are in red. Some dots may refer to more than one plot.

**Appendix S3**. Traits used for the analysis of dry calcareous grassland communities. For quantitative traits, the 5% and the 95% percentiles are shown.

| Code | Variable Name (short) | Variable Name (long) | States/ Range | Type of variable | Trait coverage % |
|---|---|---|---|---|---|
| 1 | GF_Macrophan | Growth Form – Macrophanerophyte | 0 = 452; 1 = 36 | binary | 100 |
| 2 | GF_Nanophan | Growth Form – Nanophanerophyte | 0 = 452; 1 = 36 | binary | 100 |
| 3 | GF_Hemikr | Growth Form – Hemicryptophyte | 0 = 164; 1 = 324 | binary | 100 |
| 4 | GF_Geoph | Growth Form – Geophyte | 0 = 423; 1 = 65 | binary | 100 |
| 5 | GF_Hemiphan | Growth Form – Hemiphanerophyte | 0 = 476; 1 = 12 | binary | 100 |
| 6 | GF_Theroph | Growth Form – Therophyte | 0 = 437; 1 = 51 | binary | 100 |
| 7 | GF_Pseudophan | Growth Form – Pseudophanerophyte | 0 = 484; 1 = 4 | binary | 100 |
| 8 | GF_Chamaeph | Growth Form – Chamaephyte | 0 = 458; 1 = 30 | binary | 100 |
| 9 | VP_absent | Vegetative Propagation – Absent | 0 = 279; 1 = 209 | binary | 100 |
| 10 | VP_RootShoot | Vegetative Propagation – Root shoot | 0 = 433; 1 = 55 | binary | 100 |
| 11 | VP_Runner | Vegetative Propagation – Runner | 0 = 376; 1 = 112 | binary | 100 |
| 12 | VP_Rhizome | Vegetative Propagation – Rhizome | 0 = 399; 1 = 89 | binary | 100 |
| 13 | VP_RootTuber | Vegetative Propagation – Root tuber | 0 = 473; 1 = 15 | binary | 100 |
| 14 | VP_StorageRoot | Vegetative Propagation – Storage tuber | 0 = 487; 1 = 1 | binary | 100 |
| 15 | VP_RunnerTub | Vegetative Propagation – Runner with tuberous tip | 0 = 487; 1 = 1 | binary | 100 |
| 16 | VP_Broodshoot | Vegetative Propagation – Brood shoot | 0 = 487; 1 = 1 | binary | 100 |
| 17 | VP_Fragm | Vegetative Propagation – Fragmentation | 0 = 476; 1 = 12 | binary | 100 |
| 18 | VP_Turio | Vegetative Propagation – Turio | 0 = 487; 1 = 1 | binary | 100 |
| 19 | VP_ShootTuber | Vegetative Propagation – Shoot tuber | 0 = 484; 1 = 4 | binary | 100 |
| 20 | VP_PhyllShoot | Vegetative Propagation – Phyllogenous shoot | 0 = 487; 1 = 1 | binary | 100 |
| 21 | VP_RhizPleio | Vegetative Propagation – Rhizome-like pleiocorm | 0 = 462; 1 = 26 | binary | 100 |
| 22 | VP_Bulb | Vegetative Propagation – Bulb | 0 = 485; 1 = 3 | binary | 100 |
| 23 | VP_RunnerRhiz | Vegetative Propagation – Runner-like rhizome | 0 = 485; 1 = 3 | binary | 100 |
| 24 | VP_Bulbil | Vegetative Propagation – Bulbil | 0 = 484; 1 = 4 | binary | 100 |
| 25 | FP_Beg | Flowering Phenology – Begin (month) | 3 - 7 | quantitative | 100 |
| 26 | FP_End | Flowering Phenology – End (month) | 5 - 10 | quantitative | 100 |
| 27 | FP_Dur | Flowering Phenology – Duration (month) | 2 - 5 | quantitative | 100 |
| 28 | LA | Leaf Area ($mm^2$) | 29.3 - 5960.4 | quantitative | 100 |
| 29 | SLA | Specific Leaf Area ($m^2kg^{-1}$) | 12 - 35.4 | quantitative | 100 |
| 30 | LeafCdrymass | Leaf Carbon dry mass (mg $g^{-1}$) | 419.7 - 492.4 | quantitative | 100 |
| 31 | LeafN | Leaf Nitrogen (mg $g^{-1}$) | 13.9 - 36.8 | quantitative | 100 |
| 32 | LeafP | Leaf Phosphorous (mg $g^{-1}$) | 1 - 3.6 | quantitative | 100 |
| 33 | Height | Height (m) | 0.1 - 8.9 | quantitative | 100 |
| 34 | SeedMass | Seed mass (mg) | 0 - 31.4 | quantitative | 100 |
| 35 | SeedLength | Seed length (mm) | 0.6 - 8.8 | quantitative | 100 |
| 36 | LDMC | Leaf dry Matter Content (g $g^{-1}$) | 0.1 - 0.4 | quantitative | 100 |
| 37 | LeafNarea | Leaf Nitrogen per area (g $m^{-2}$) | 0.7 - 2 | quantitative | 100 |
| 38 | Leaf N:P ratio | Leaf N:P ratio (g $g^{-1}$) | 6.4 - 19.6 | quantitative | 100 |
| 39 | Leaf d15N | Leaf $\delta^{15}N$ (ppm) | 0.4 - 5.6 | quantitative | 100 |
| 40 | SeedNum | Seed number per reproductive unit | 12.6 - 39957.5 | quantitative | 100 |
| 41 | LeafFreshMass | Leaf fresh mass (g) | 0 - 1.7 | quantitative | 100 |
| 42 | DispUnitLength | Dispersal unit length (mm) | 0.6 - 11.6 | quantitative | 100 |
| 43 | LeafPersistence | Leaf persistence | Spring-green, Summer-green, Winter-green, Evergreen | nominal | 99.8 |
| 44 | Pollination | Pollination | Nectar/Honey/Insects, Pollen, Wind | nominal | 96.5 |
| 45 | ReprStrat | Reproduction strategy | Seed/Spores, Vegetative | nominal | 100 |
| 46 | Sclerophylly | Sclerophylly | 1, 1.5, 2, 2.5, 3, 3.5, 4 | ordinal | 99.7 |
| 47 | Leaf_Succulent | Leaf succulence | 0 = 473; 1 = 6 | binary | 99.9 |
| 48 | LifeSpan | Lifespan | 1, 1.5, 2, 2.5, 3 | ordinal | 100 |
| 49 | Rosette | Rosette | 0, 0.5, 1 | ordinal | 100 |

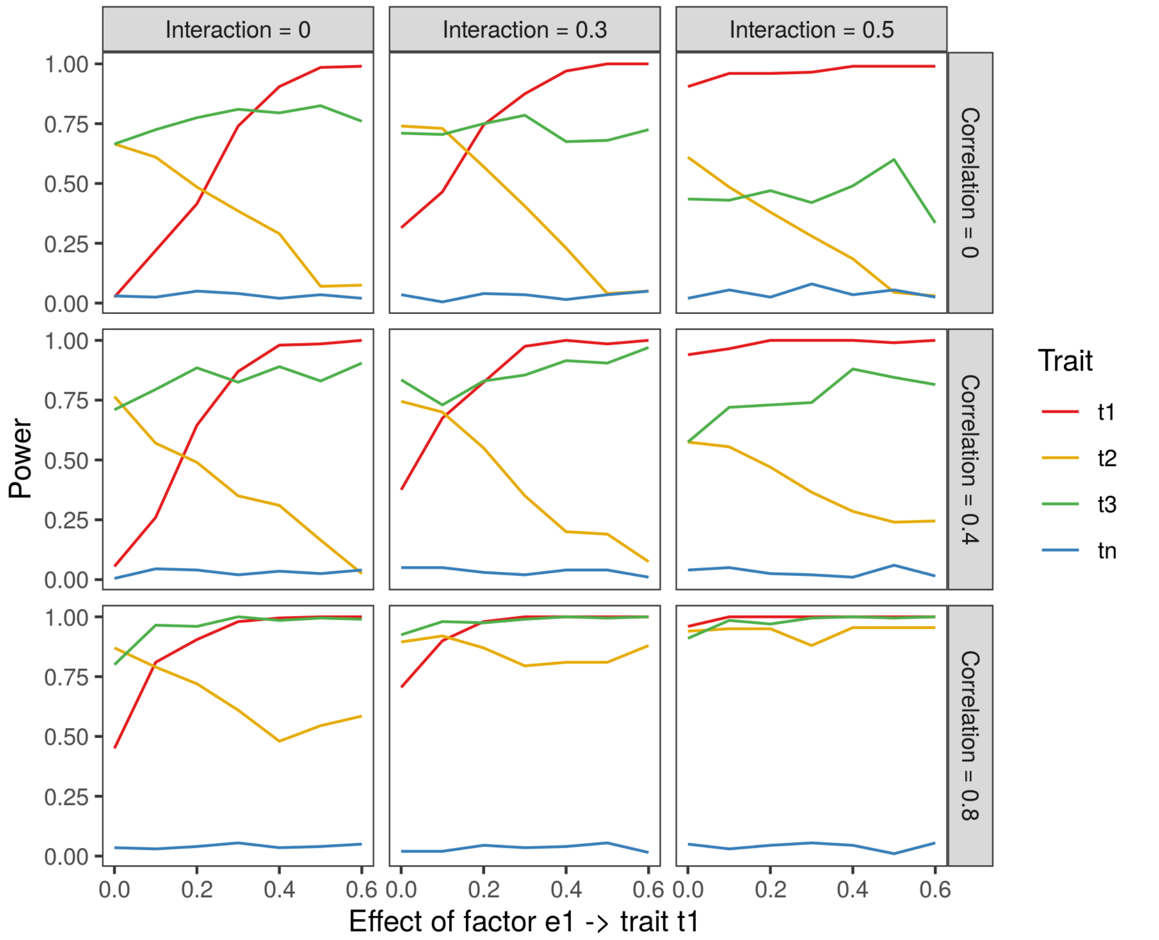


**Appendix S4**. Simulated-data power profiles of the Rd matrix correlation r(**XY**) between community distances based on trait-based fuzzy-weighted (**X**) and Beals-smoothed (**Y**) species composition for metacommunities. A total of 100 metacommunities was simulated under Scenario 2 by a stochastic individual-based model with increasing strength of factor effect $e_1$ on trait $t_1$, using linear parameters for this main effect ranging from 0 to 0.6 (horizontal axes), a fixed strength of 0.3 for factor effects $e_2$ -> $t_2$ and $e_1$ -> $t_3$, and no factor effect on trait $t_n$. Also, the pair-wise correlations between traits $t_1$, $t_2$ and $t_3$ ranged from 0 to 0.8 (upper to bottom panel), and the parameter for factor interaction $e_1$ x $e_2$ ranged from 0 to 0.5 (left to right panels). Factor interaction effects were zero. Power (vertical axis) is the proportion of simulated metacommunities for which the P-value for r(**XY**) found by permutation was not larger than a threshold of 0.05. The graphs show only traits considered individually to define fuzzy-weighted species composition.

**Appendix S5**. Numerical results for best solutions shown in Figure 5. Trait codes are shown in Appendix S3.

| Trait combination | Num. traits (tier) | r(XY) observed | Expected r(XY) under H0 | Lower limit | Upper limit | p-value | Optimal at tier? |
|---|---|---|---|---|---|---|---|
| 46_27_17_33_29_5_2 | 7 | 0.466 | 0.155 | 0.444 | 0.509 | 0.001 | FALSE |
| 46_27_17_33_29_5 | 6 | 0.480 | 0.236 | 0.458 | 0.527 | 0.001 | FALSE |
| 46_27_17_33_29 | 5 | 0.474 | 0.202 | 0.455 | 0.521 | 0.001 | TRUE |
| 46_27_17_33 | 4 | 0.458 | 0.203 | 0.438 | 0.505 | 0.001 | FALSE |
| 46_27_33 | 3 | 0.429 | 0.150 | 0.412 | 0.476 | 0.001 | FALSE |
| 46_27 | 2 | 0.399 | 0.140 | 0.377 | 0.449 | 0.001 | TRUE |
| 46 | 1 | 0.317 | 0.119 | 0.286 | 0.367 | 0.003 | TRUE |
| 33 | 1 | 0.267 | 0.110 | 0.247 | 0.319 | 0.012 | FALSE |
| 29 | 1 | 0.251 | 0.094 | 0.229 | 0.294 | 0.022 | FALSE |
| 2 | 1 | 0.250 | 0.044 | 0.218 | 0.294 | 0.019 | FALSE |
| 27 | 1 | 0.237 | 0.110 | 0.212 | 0.285 | 0.032 | FALSE |
| 5 | 1 | 0.210 | 0.037 | 0.174 | 0.270 | 0.033 | FALSE |
| 17 | 1 | 0.207 | 0.217 | 0.170 | 0.272 | 0.031 | FALSE |

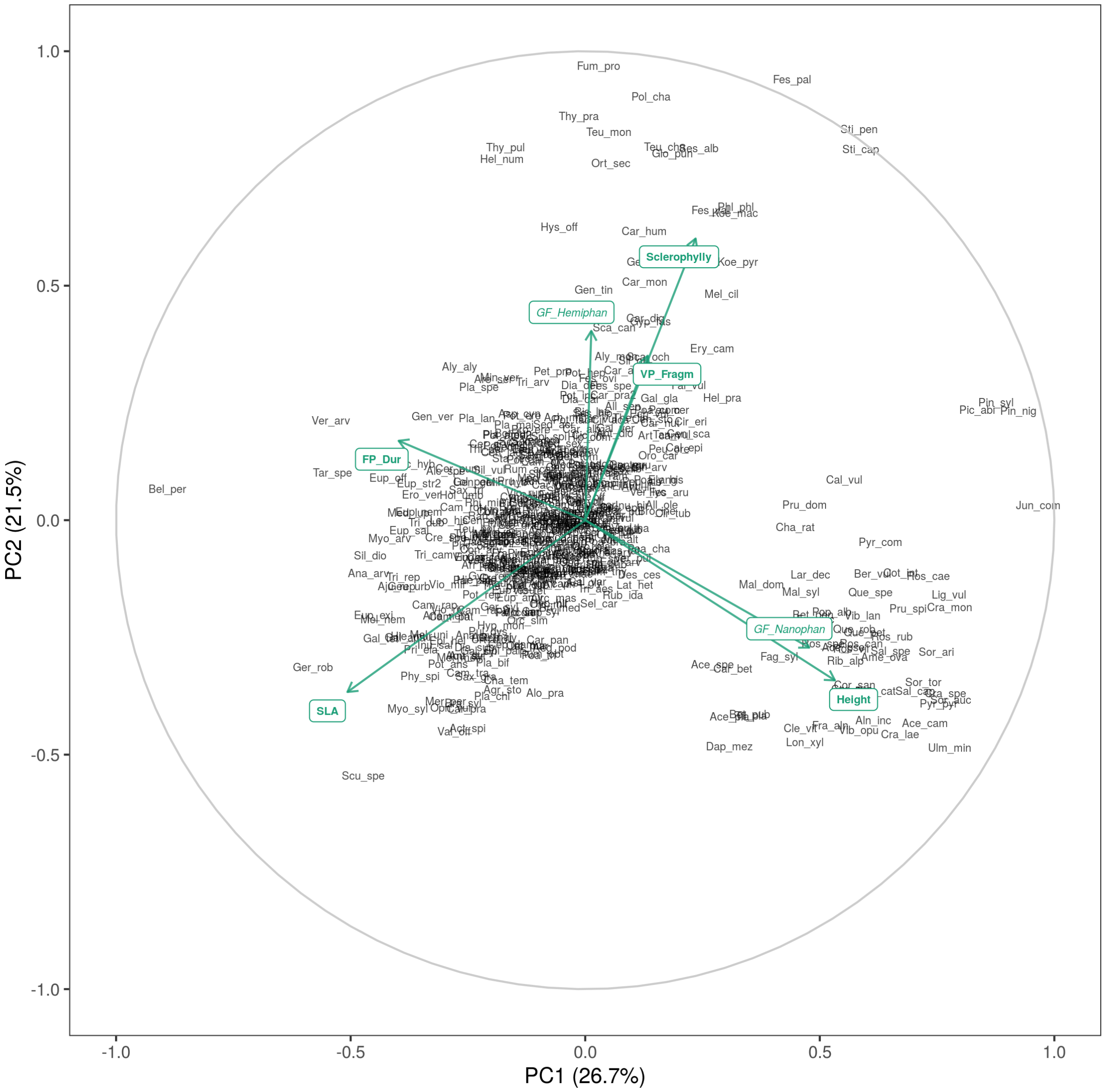


**Appendix S6.** Principal components analysis (PCA) of species on the basis of the matrix of 488 species by the 7 traits that were relevant in community assembly (Fig. 5). The species were found in 565 plots of calcareous dry grassland in Germany. See Appendix S13 for species labels. The PCA was based on pairwise trait correlations. The traits are projected on the ordination space according to their Pearson correlation with the ordination axes.

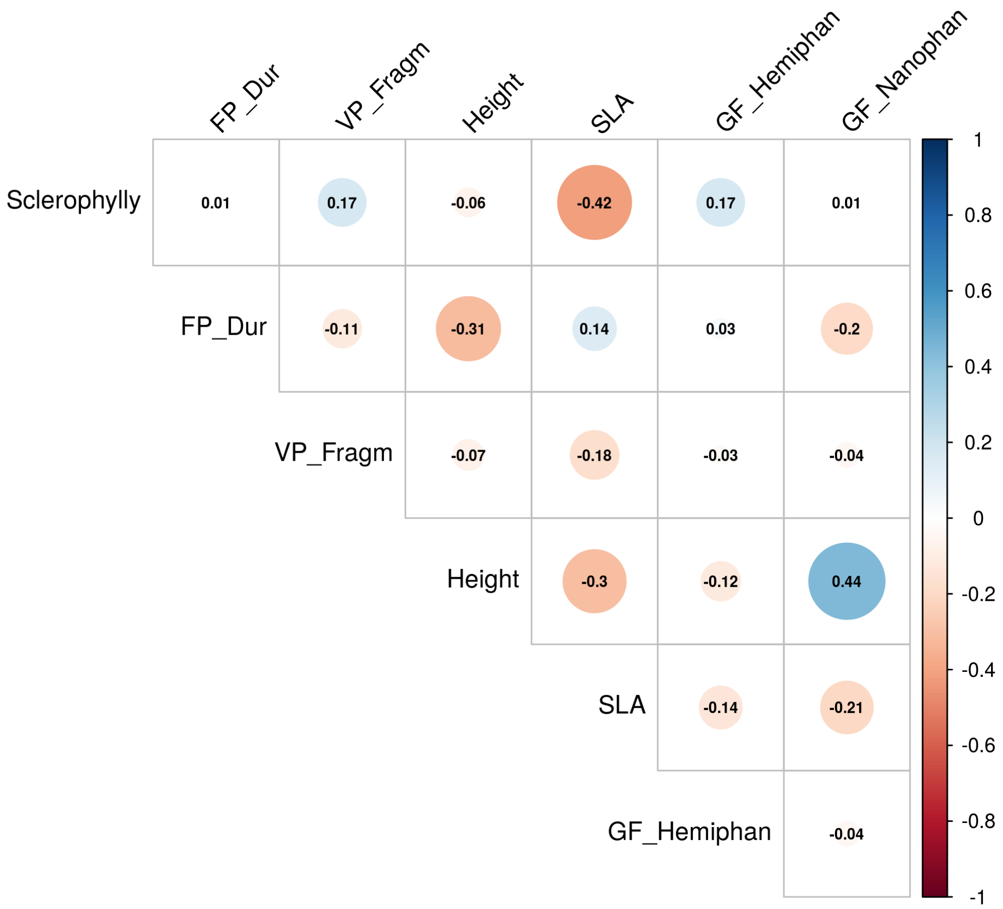


**Appendix S7**. Pearson correlation matrix between traits at species level, based on the matrix of 488 species by the 7 traits found relevant in community assembly (Fig. 5). The species were the ones occurring in 565 plots of calcareous dry grassland in Germany.

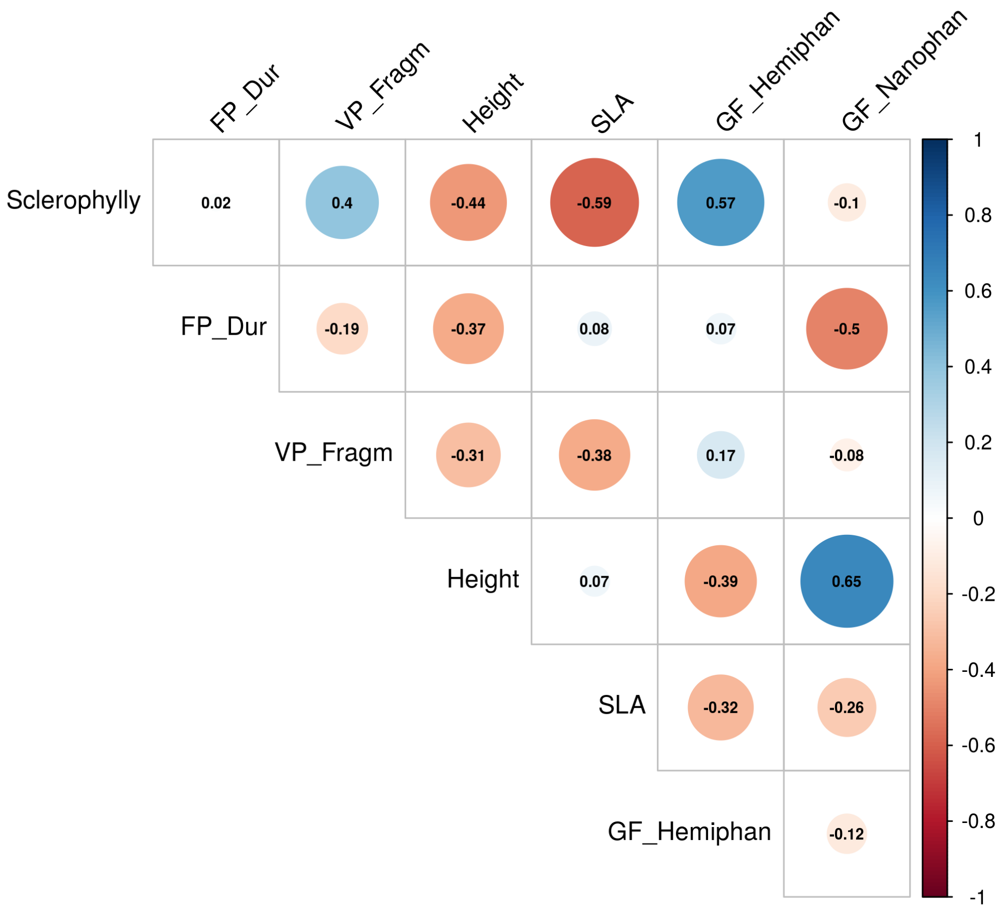


**Appendix S8.** Pearson correlation matrix between community weighted means (CWM) for the 7 quantitative traits found relevant in community assembly (Fig. 5), considering 565 plots of calcareous dry grassland in Germany.

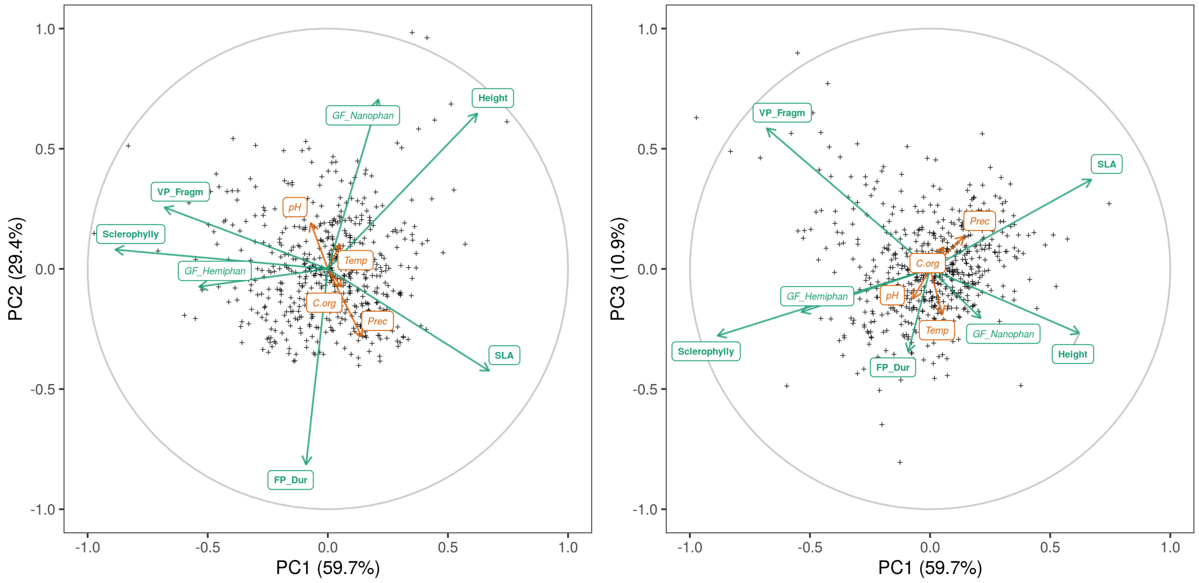


**Appendix S9**. Principal components analysis (PCA) of dry grassland plots (cross symbols) based on the fuzzy-weighted composition (**X** matrix) defined by the five optimal traits identified in Fig. 5. The CWMs for the traits with a significant Rd matrix correlation r(**XY**) in Fig. 4, in green, and four environmental variables, in orange, are projected on the ordination space according to their Pearson correlations with the PCA axes. The five traits identified in Fig. 5 as the best combination of traits are shown in bold green fonts.

**Appendix S10.** Correlation (Pearson) coefficients between principal components of Beals' smoothed composition (**Y** matrix) and community-weighted mean trait values (CWMs), shown in green, environmental variables, shown in orange, and principal components of the fuzzy-weighted composition (**X** matrix), shown in purple. The five traits identified in Fig. 5 as the best combination of traits for predicting fuzzy-weighted species composition related to species co-occurrences are shown in bold green fonts.

| **CWM trait, environmental factor, or PC fuzzy-weighted composition** | **PC1** | **PC2** | **PC3** | **PC4** |
|---|---|---|---|---|
| FP_Dur | -0.450 | 0.243 | -0.073 | 0.169 |
| GF_Hemiphan | 0.314 | 0.352 | -0.322 | 0.233 |
| GF_Nanophan | 0.332 | -0.399 | -0.026 | -0.266 |
| Height | 0.120 | -0.353 | 0.386 | -0.372 |
| Sclerophylly | 0.501 | 0.284 | -0.403 | 0.349 |
| SLA | -0.399 | -0.088 | 0.569 | -0.085 |
| VP_Fragm | 0.322 | 0.301 | -0.371 | -0.229 |
| Temp | 0.178 | -0.043 | 0.153 | 0.165 |
| Prec | -0.386 | 0.025 | -0.052 | -0.261 |
| pH | 0.247 | -0.087 | -0.005 | 0.152 |
| C.org | -0.131 | 0.096 | 0.062 | -0.171 |
| FW-PC1 | -0.407 | -0.369 | 0.582 | -0.209 |
| FW-PC2 | 0.553 | -0.213 | 0.002 | -0.275 |
| FW-PC3 | -0.089 | 0.070 | 0.023 | -0.277 |

Temp: mean annual temperature, Prec: mean annual precipitation, from Chelsa. pH: soil pH, C.org: soil organic carbon. From SOILGRIDS.

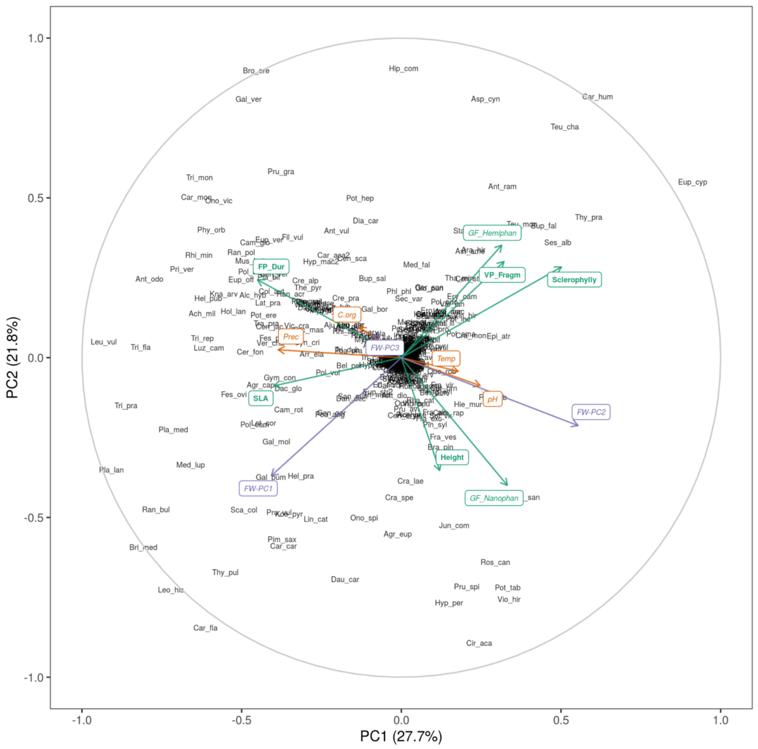


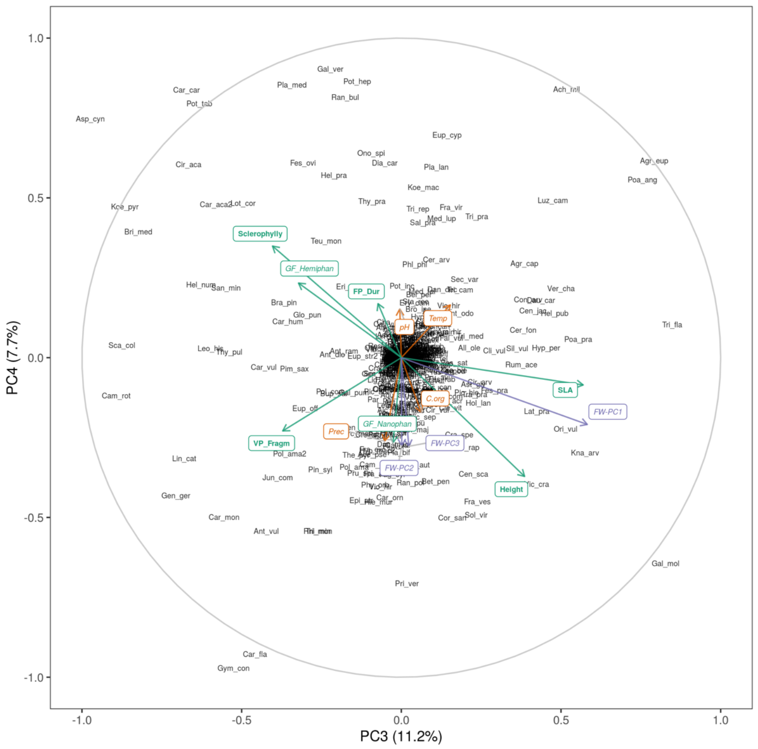


**Appendix S11**. Principal components analysis (PCA) of dry grassland plots described by the Beals' smoothed composition (**Y** matrix). This is the same ordination of Fig. 6, but the species eigenvectors are plotted (see Appendix S13 for species labels). The CWMs for the traits with a significant Rd matrix correlation r(**XY**) in Fig. 4, in green, are projected on the ordination space according to their Pearson correlations with the PCA axes (see correlations in Appendix S10). The five traits identified in Fig. 5 as the best combination of traits are shown in bold green fonts.

**Appendix S12.** Identification numbers of the 565 dry calcareous grassland relevés in Germany extracted from the GVRD database.

Relevé numbers: 919 1899 3871 3886 3900 3925 3940 3943 3949 3950 3952 3961 3965 3971 3984 3988 3990 3992 3993 4018 4034 4444 4455 4458 4461 4467 4489 4503 4540 4542 4562 4569 4588 4593 4597 4617 4620 4621 4632 4638 4664 4670 4671 4674 8031 8048 8053 8073 8084 8098 8111 8123 8131 8374 8376 10732 10737 10749 11846 11864 11869 14636 14639 14643 14650 17227 17231 17235 17236 17246 17247 17358 17362 17373 17376 17413 17415 17419 17422 17425 17431 17433 17442 17445 25695 25697 25699 25706 25717 25728 25741 25743 25746 25804 25805 25823 25830 25843 25853 25858 25866 25869 25871 25875 25878 25880 25886 25903 25909 25911 25912 25951 25952 25960 25964 25966 25987 26001 26017 26029 26070 26076 26086 26146 26165 26166 26167 26187 26188 26192 26223 26224 26235 26237 26269 26294 26300 26303 26304 26311 26312 26315 26328 26344 26357 26367 26369 26384 26389 30673 35185 35190 35191 35196 35201 35202 35981 35987 35993 35999 36010 36707 36720 36724 36727 36737 36739 36760 36766 36769 36771 36774 36796 36798 36828 36837 36841 36842 36860 36863 36868 36880 36885 36887 36899 36902 36911 36920 36922 36927 37153 37160 37166 37209 37212 37213 37241 37247 37250 37251 37259 37263 37264 37280 37330 37344 37355 37378 37402 37404 37416 37466 37472 37513 37559 37571 37573 37575 37601 37629 37636 37643 37658 37660 37669 37671 37685 37694 37702 37710 37714 37719 37749 37761 37762 37787 37813 37839 37873 37917 37919 37922 37926 37944 37946 37976 37980 37994 38001 38023 38042 38091 38093 38108 38109 38131 38134 38147 38152 38167 38175 38176 38193 38314 38315 38318 38361 38365 38391 38401 38408 38421 38425 38438 38442 38471 38480 38481 38488 38496 38507 38509 38544 38562 38595 38650 38657 38669 38694 38729 38734 38755 38759 38789 38790 38806 38807 38827 38832 38834 38839 38864 38883 38884 38887 38890 38892 38897 38924 38926 38927 38933 38985 38998 38999 39007 39008 39012 39014 39038 39069 39086 39099 39103 39119 39120 39131 39132 39171 39198 39199 39200 39201 39206 39213 39217 39223 39227 39280 39286 39297 39417 39422 39438 39441 39442 39602 39603 39605 39608 39753 39755 39759 39770 39773 39779 39781 39809 39812 39814 39816 39836 39996 39998 40001 40018 40021 40023 40030 40031 40037 40039 40044 40051 40066 40082 40106 40107 40110 40127 40131 40132 40149 40161 40167 40175 40189 40212 40240 40242 40253 40262 40264 40265 40275 40284 40308 40316 40317 40329 40332 40339 40344 40350 40371 40373 40460 40473 40477 40481 40485 40495 40496 40539 40541 40560 40569 40583 40586 40606 40636 40652 40739 40794 40805 40810 40813 40820 40828 40829 40832 40847 40856 40865 40866 40875 40909 40910 40911 40916 40932 40943 40950 40984 41002 41003 41011 41031 41442 41443 41452 42265 42275 42316 42320 42330 42334 42341 42402 42417 42418 42420 42432 42451 42586 42588 42599 42600 42605 42606 42778 42795 42806 42816 43122 43134 43138 43177 43670 43699 43704 43720 43724 43725 43734 44000 44001 44122 44151 44159 44169 44195 44199 44204 44213 44273 44285 44291 44300 44318 44319 44326 44335 44347 44349 44360 44364 44367 44369 44379 44389 44396 44399 44401 44405 44406 44429 44435 44442 44445 44486 44489 44645 44646 44699 44700 44706 44715 44723 44747 44751 44804 44815 44821 44827 44836 45006 45035 51609 55260 55266 55278 55281 55510 55511 55520 55529 55530 57336 57337 57342 58145 58153 58155 58916 58917 58925 58944 58946 58959 60693 60941 60942 60960 62090

**Appendix S13.** Species (488) found in 565 dry calcareous grassland relevés extracted from the GVRD database.

| Label | Species | Label | Species | Label | Species |
|---|---|---|---|---|---|
| Ace_cam | Acer campestre | Bri_med | Briza media | Cli_vul | Clinopodium vulgare |
| Ace_pla | Acer platanoides | Bro_ere | Bromus erectus | Col_aut | Colchicum autumnale |
| Ace_pse | Acer pseudoplatanus | Bro_hor | Bromus hordeaceus | Con_maj | Convallaria majalis |
| Ace_spe | Acer spec | Bro_ine | Bromus inermis | Con_arv | Convolvulus arvensis |
| Ach_mil | Achillea millefolium | Bro_ste | Bromus sterilis | Cor_san | Cornus sanguinea |
| Act_spi | Actaea spicata | Bun_bul | Bunium bulbocastanum | Cor_vag | Coronilla vaginalis |
| Ado_ver | Adonis vernalis | Bup_sal | Buphthalmum salicifolium | Cor_ave | Corylus avellana |
| Aeg_pod | Aegopodium podagraria | Bup_fal | Bupleurum falcatum | Cot_int | Cotoneaster integerrimus |
| Agr_eup | Agrimonia eupatoria | Cal_epi | Calamagrostis epigejos | Cra_lae | Crataegus laevigata |
| Agr_cap | Agrostis capillaris | Cal_var | Calamagrostis varia | Cra_mon | Crataegus monogyna |
| Agr_spe | Agrostis spec | Cal_vul | Calluna vulgaris | Cra_spe | Crataegus spec |
| Agr_sto | Agrostis stolonifera agg | Cam_glo | Campanula glomerata | Cre_alp | Crepis alpestris |
| Air_car | Aira caryophyllea | Cam_pat | Campanula patula | Cre_bie | Crepis biennis |
| Aju_gen | Ajuga genevensis | Cam_per | Campanula persicifolia | Cre_mol | Crepis mollis |
| Aju_rep | Ajuga reptans | Cam_rap | Campanula rapunculoides | Cre_pra | Crepis praemorsa |
| Alc_hyb | Alchemilla hybrida agg | Cam_rap2 | Campanula rapunculus | Cre_spe | Crepis spec |
| Alc_spe | Alchemilla spec | Cam_rot | Campanula rotundifolia | Cru_lae | Cruciata laevipes |
| All_ole | Allium oleraceum | Cam_tra | Campanula trachelium | Cus_epi | Cuscuta epithymum |
| All_sen | Allium senescens | Car_pra | Cardamine pratensis | Cyn_dac | Cynodon dactylon |
| All_vin | Allium vineale | Car_aca | Carduus acanthoides | Cyn_off | Cynoglossum officinale |
| Aln_inc | Alnus incana | Car_nut | Carduus nutans | Cyn_cri | Cynosurus cristatus |
| Alo_pra | Alopecurus pratensis | Car_alb | Carex alba | Dac_glo | Dactylis glomerata |
| Aly_aly | Alyssum alyssoides | Car_car | Carex caryophyllea | Dac_mac | Dactylorhiza maculata |
| Aly_mon | Alyssum montanum | Car_dig | Carex digitata | Dan_dec | Danthonia decumbens |
| Ame_ova | Amelanchier ovalis | Car_fla | Carex flacca | Dap_mez | Daphne mezereum |
| Ana_pyr | Anacamptis pyramidalis | Car_hir | Carex hirta | Dau_car | Daucus carota |
| Ana_arv | Anagallis arvensis | Car_hum | Carex humilis | Des_ces | Deschampsia cespitosa |
| Ane_nar | Anemone narcissiflora | Car_mon | Carex montana | Des_fle | Deschampsia flexuosa |
| Ane_nem | Anemone nemorosa | Car_mur | Carex muricata agg | Dia_arm | Dianthus armeria |
| Ane_syl | Anemone sylvestris | Car_orn | Carex ornithopoda | Dia_car | Dianthus carthusianorum |
| Ant_dio | Antennaria dioica | Car_pan | Carex panicea | Dia_del | Dianthus deltoides |
| Ant_lil | Anthericum liliago | Car_pil | Carex pilulifera | Dia_sup | Dianthus superbus |
| Ant_ram | Anthericum ramosum | Car_pra2 | Carex praecox | Ech_vul | Echium vulgare |
| Ant_odo | Anthoxanthum odoratum | Car_pul | Carex pulicaris | Ely_his | Elymus hispidus |
| Ant_syl | Anthriscus sylvestris | Car_syl | Carex sylvatica | Ely_rep | Elymus repens |
| Ant_vul | Anthyllis vulneraria | Car_aca2 | Carlina acaulis | Epi_atr | Epipactis atrorubens |
| Aqu_atr | Aquilegia atrata | Car_vul | Carlina vulgaris | Epi_hel | Epipactis helleborine |
| Aqu_vul | Aquilegia vulgaris | Car_bet | Carpinus betulus | Epi_pal | Epipactis palustris |
| Ara_hir | Arabis hirsuta | Car_car2 | Carum carvi | Equ_arv | Equisetum arvense |
| Ara_sag | Arabis sagittata | Cau_pla | Caucalis platycarpos | Eri_acr | Erigeron acris |
| Are_ser | Arenaria serpyllifolia agg | Cen_jac | Centaurea jacea | Ero_ver | Erophila verna |
| Arr_ela | Arrhenatherum elatius | Cen_sca | Centaurea scabiosa | Ery_cam | Eryngium campestre |
| Art_cam | Artemisia campestris | Cen_sto | Centaurea stoebe | Eup_amy | Euphorbia amygdaloides |
| Asp_cyn | Asperula cynanchica | Cen_ery | Centaurium erythraea | Eup_cyp | Euphorbia cyparissias |
| Asp_tin | Asperula tinctoria | Cep_dam | Cephalanthera damasonium | Eup_esu | Euphorbia esula |
| Asp_tri | Asplenium trichomanes | Cep_rub | Cephalanthera rubra | Eup_exi | Euphorbia exigua |
| Ast_ame | Aster amellus | Cer_arv | Cerastium arvense | Eup_seg | Euphorbia seguieriana |
| Ast_dan | Astragalus danicus | Cer_fon | Cerastium fontanum agg | Eup_str | Euphorbia stricta |
| Ast_gly | Astragalus glycyphyllos | Cer_pum | Cerastium pumilum agg | Eup_ver | Euphorbia verrucosa |
| Ast_maj | Astrantia major | Cer_tom | Cerastium tomentosum | Eup_nem | Euphrasia nemorosa agg |
| Bel_per | Bellis perennis | Cha_tem | Chaerophyllum temulum | Eup_off | Euphrasia officinalis |
| Ber_vul | Berberis vulgaris | Cha_rat | Chamaecytisus ratisbonensis | Eup_sal | Euphrasia salisburgensis |
| Bet_pen | Betula pendula | Cic_int | Cichorium intybus | Eup_str2 | Euphrasia stricta |
| Bet_pub | Betula pubescens | Cir_aca | Cirsium acaule | Fag_syl | Fagus sylvatica |
| Bis_lae | Biscutella laevigata | Cir_arv | Cirsium arvense | Fal_vul | Falcaria vulgaris |
| Bot_isc | Bothriochloa ischaemum | Cir_eri | Cirsium eriophorum | Fal_con | Fallopia convolvulus |
| Bot_lun | Botrychium lunaria | Cir_tub | Cirsium tuberosum | Fes_aru | Festuca arundinacea |
| Bra_pin | Brachypodium pinnatum | Cir_vul | Cirsium vulgare | Fes_het | Festuca heterophylla |
| Bra_syl | Brachypodium sylvaticum | Cle_vit | Clematis vitalba | Fes_ovi | Festuca ovina agg |

| Label | Species | Label | Species | Label | Species |
|---|---|---|---|---|---|
| Fes_pal | Festuca pallens | Hyp_mac2 | Hypochaeris maculata | Orc_sim | Orchis simia |
| Fes_pra | Festuca pratensis | Hyp_rad | Hypochaeris radicata | Ori_vul | Origanum vulgare |
| Fes_rub | Festuca rubra agg | Hys_off | Hyssopus officinalis | Oro_car | Orobanche caryophyllacea |
| Fes_spe | Festuca spec | Inu_hir | Inula hirta | Oro_ela | Orobanche elatior |
| Fes_val | Festuca valesiaca | Inu_sal | Inula salicina | Oro_lut | Orobanche lutea |
| Fil_vul | Filipendula vulgaris | Jun_com | Juniperus communis | Oro_min | Orobanche minor |
| Fra_spe | Fragaria spec | Kna_arv | Knautia arvensis | Oro_teu | Orobanche teucrii |
| Fra_ves | Fragaria vesca | Kna_dip | Knautia dipsacifolia | Ort_sec | Orthilia secunda |
| Fra_vir | Fragaria viridis | Koe_mac | Koeleria macrantha | Par_pal | Parnassia palustris |
| Fra_aln | Frangula alnus | Koe_pyr | Koeleria pyramidata | Pas_sat | Pastinaca sativa |
| Fra_exc | Fraxinus excelsior | Lac_per | Lactuca perennis | Pet_pro | Petrorhagia prolifera |
| Fum_pro | Fumana procumbens | Lar_dec | Larix decidua | Peu_als | Peucedanum alsaticum |
| Gal_tet | Galeopsis tetrahit | Las_lat | Laserpitium latifolium | Peu_cer | Peucedanum cervaria |
| Gal_apa | Galium aparine | Lat_het | Lathyrus heterophyllus | Peu_ore | Peucedanum oreoselinum |
| Gal_bor | Galium boreale | Lat_lin | Lathyrus linifolius | Phl_ber | Phleum bertolonii |
| Gal_gla | Galium glaucum | Lat_pra | Lathyrus pratensis | Phl_phl | Phleum phleoides |
| Gal_mol | Galium mollugo agg | Lat_spe | Lathyrus spec | Phl_pra | Phleum pratense |
| Gal_pum | Galium pumilum | Lat_tub | Lathyrus tuberosus | Phy_orb | Phyteuma orbiculare |
| Gal_syl | Galium sylvaticum | Leo_his | Leontodon hispidus | Phy_spi | Phyteuma spicatum |
| Gal_ver | Galium verum | Leu_vul | Leucanthemum vulgare agg | Pic_abi | Picea abies |
| Gen_pil | Genista pilosa | Lig_vul | Ligustrum vulgare | Pic_hie | Picris hieracioides |
| Gen_tin | Genista tinctoria | Lin_vul | Linaria vulgaris | Pim_maj | Pimpinella major |
| Gen_cru | Gentiana cruciata | Lin_cat | Linum catharticum | Pim_sax | Pimpinella saxifraga |
| Gen_lut | Gentiana lutea | Lin_leo | Linum leonii | Pin_nig | Pinus nigra |
| Gen_ver | Gentiana verna | Lin_ten | Linum tenuifolium | Pin_syl | Pinus sylvestris |
| Gen_ger | Gentianella germanica | Lit_off | Lithospermum officinale | Pla_lan | Plantago lanceolata |
| Ger_col | Geranium columbinum | Lol_per | Lolium perenne | Pla_maj | Plantago major |
| Ger_rob | Geranium robertianum | Lon_xyl | Lonicera xylosteum | Pla_med | Plantago media |
| Ger_san | Geranium sanguineum | Lot_cor | Lotus corniculatus | Pla_spe | Plantago spec |
| Ger_syl | Geranium sylvaticum | Luz_cam | Luzula campestris | Pla_bif | Platanthera bifolia |
| Geu_riv | Geum rivale | Luz_luz | Luzula luzuloides | Pla_chl | Platanthera chlorantha |
| Geu_urb | Geum urbanum | Luz_mul | Luzula multiflora | Poa_ang | Poa angustifolia |
| Glo_pun | Globularia punctata | Mal_dom | Malus domestica | Poa_cha | Poa chaixii |
| Goo_rep | Goodyera repens | Mal_syl | Malus sylvestris | Poa_com | Poa compressa |
| Gym_con | Gymnadenia conopsea | Med_fal | Medicago falcata | Poa_nem | Poa nemoralis |
| Gym_odo | Gymnadenia odoratissima | Med_lup | Medicago lupulina | Poa_pra | Poa pratensis agg |
| Gyp_fas | Gypsophila fastigiata | Med_sat | Medicago sativa agg | Poa_tri | Poa trivialis |
| Gyp_rep | Gypsophila repens | Mel_arv | Melampyrum arvense | Pol_ama | Polygala amara agg |
| Hel_num | Helianthemum nummularium | Mel_cri | Melampyrum cristatum | Pol_ama2 | Polygala amarella |
| Hel_pra | Helictotrichon pratense | Mel_nem | Melampyrum nemorosum | Pol_cal | Polygala calcarea |
| Hel_pub | Helictotrichon pubescens | Mel_cil | Melica ciliata | Pol_cha | Polygala chamaebuxus |
| Hep_nob | Hepatica nobilis | Mel_uni | Melica uniflora | Pol_com | Polygala comosa |
| Her_sph | Heracleum sphondylium | Mel_alb | Melilotus albus | Pol_vul | Polygala vulgaris |
| Her_mon | Herminium monorchis | Mel_alt | Melilotus altissimus | Pol_odo | Polygonatum odoratum |
| Hie_bif | Hieracium bifidum | Mel_off | Melilotus officinalis | Pop_alb | Populus alba |
| Hie_cae | Hieracium caesium | Mel_mel | Melittis melissophyllum | Pop_tre | Populus tremula |
| Hie_cym | Hieracium cymosum | Mer_per | Mercurialis perennis | Pot_ans | Potentilla anserina |
| Hie_gla | Hieracium glaucinum | Min_hyb | Minuartia hybrida | Pot_ere | Potentilla erecta |
| Hie_lae | Hieracium laevigatum | Min_ver | Minuartia verna | Pot_hep | Potentilla heptaphylla |
| Hie_mur | Hieracium murorum | Mus_bot | Muscari botryoides | Pot_inc | Potentilla incana |
| Hie_sab | Hieracium sabaudum | Myo_arv | Myosotis arvensis | Pot_rep | Potentilla reptans |
| Hie_spe | Hieracium spec | Myo_syl | Myosotis sylvatica | Pot_tab | Potentilla tabernaemontani |
| Hie_umb | Hieracium umbellatum | Odo_ver | Odontites vernus agg | Pri_ela | Primula elatior |
| Him_hir | Himantoglossum hircinum | Ono_are | Onobrychis arenaria | Pri_ver | Primula veris |
| Hip_com | Hippocrepis comosa | Ono_vic | Onobrychis viciifolia | Pru_gra | Prunella grandiflora |
| Hol_lan | Holcus lanatus | Ono_spi | Ononis spinosa agg | Pru_lac | Prunella laciniata |
| Hol_umb | Holosteum umbellatum | Oph_vul | Ophioglossum vulgatum | Pru_vul | Prunella vulgaris |
| Hyp_ele | Hypericum elegans | Oph_api | Ophrys apifera | Pru_avi | Prunus avium |
| Hyp_hir | Hypericum hirsutum | Oph_ins | Ophrys insectifera | Pru_dom | Prunus domestica |
| Hyp_mac | Hypericum maculatum | Orc_mas | Orchis mascula | Pru_spi | Prunus spinosa |
| Hyp_mon | Hypericum montanum | Orc_mil | Orchis militaris | Pul_dys | Pulicaria dysenterica |
| Hyp_per | Hypericum perforatum | Orc_pur | Orchis purpurea | Pyr_com | Pyrus communis |

| Label | Species |
|---|---|
| Pyr_pyr | Pyrus pyraster |
| Que_pet | Quercus petraea |
| Que_rob | Quercus robur |
| Que_spe | Quercus spec |
| Ran_acr | Ranunculus acris |
| Ran_bul | Ranunculus bulbosus |
| Ran_pol | Ranunculus polyanthemos agg |
| Res_lut | Reseda lutea |
| Rha_cat | Rhamnus cathartica |
| Rhi_ale | Rhinanthus alectorolophus agg |
| Rhi_gla | Rhinanthus glacialis |
| Rhi_min | Rhinanthus minor |
| Rib_alp | Ribes alpinum |
| Ros_cae | Rosa caesia |
| Ros_can | Rosa canina |
| Ros_rub | Rosa rubiginosa agg |
| Ros_spe | Rosa spec |
| Ros_spi | Rosa spinosissima |
| Ros_vil | Rosa villosa agg |
| Rub_cae | Rubus caesius |
| Rub_ida | Rubus idaeus |
| Rub_sax | Rubus saxatilis |
| Rub_spe | Rubus spec |
| Rum_ace | Rumex acetosa |
| Rum_ace2 | Rumex acetosella |
| Rum_obt | Rumex obtusifolius |
| Rum_thy | Rumex thyrsiflorus |
| Sal_cap | Salix caprea |
| Sal_spe | Salix spec |
| Sal_pra | Salvia pratensis |
| Sal_ver | Salvia verticillata |
| San_min | Sanguisorba minor |
| Sax_gra | Saxifraga granulata |
| Sax_ros | Saxifraga rosacea |
| Sax_tri | Saxifraga tridactylites |
| Sca_can | Scabiosa canescens |
| Sca_col | Scabiosa columbaria |
| Sca_och | Scabiosa ochroleuca |
| Scu_spe | Scutellaria spec |
| Sec_var | Securigera varia |
| Sed_acr | Sedum acre |
| Sed_alb | Sedum album |
| Sed_max | Sedum maximum |
| Sed_sex | Sedum sexangulare |
| Sed_tel | Sedum telephium agg |
| Sel_car | Selinum carvifolia |
| Ser_tin | Serratula tinctoria |
| Ses_ann | Seseli annuum |
| Ses_hip | Seseli hippomarathrum |
| Ses_lib | Seseli libanotis |
| Ses_alb | Sesleria albicans |
| Sil_sil | Silaum silaus |
| Sil_dio | Silene dioica |
| Sil_nut | Silene nutans |
| Sil_oti | Silene otites |
| Sil_vul | Silene vulgaris |
| Sol_vir | Solidago virgaurea |
| Sor_ari | Sorbus aria |
| Sor_auc | Sorbus aucuparia |
| Sor_tor | Sorbus torminalis |
| Spi_spi | Spiranthes spiralis |
| Sta_rec | Stachys recta |
| Ste_gra | Stellaria graminea |
| Sti_cap | Stipa capillata |
| Sti_pen | Stipa pennata |
| Suc_pra | Succisa pratensis |
| Tan_cor | Tanacetum corymbosum |
| Tan_vul | Tanacetum vulgare |
| Tar_spe | Taraxacum spec |
| Tet_mar | Tetragonolobus maritimus |
| Teu_bot | Teucrium botrys |
| Teu_cha | Teucrium chamaedrys |
| Teu_mon | Teucrium montanum |
| Tha_min | Thalictrum minus |
| Tha_sim | Thalictrum simplex |
| The_bav | Thesium bavarum |
| The_lin | Thesium linophyllon |
| The_pyr | Thesium pyrenaicum |
| Thl_per | Thlaspi perfoliatum |
| Thy_pra | Thymus praecox |
| Thy_pul | Thymus pulegioides agg |
| Til_pla | Tilia platyphyllos |
| Tor_jap | Torilis japonica |
| Tra_pra | Tragopogon pratensis |
| Tra_glo | Traunsteinera globosa |
| Tri_alp | Trifolium alpestre |
| Tri_arv | Trifolium arvense |
| Tri_cam | Trifolium campestre |
| Tri_dub | Trifolium dubium |
| Tri_hyb | Trifolium hybridum |
| Tri_med | Trifolium medium |
| Tri_mon | Trifolium montanum |
| Tri_och | Trifolium ochroleucon |
| Tri_pra | Trifolium pratense |
| Tri_rep | Trifolium repens |
| Tri_rub | Trifolium rubens |
| Tri_fla | Trisetum flavescens |
| Tri_aes | Triticum aestivum |
| Tro_eur | Trollius europaeus |
| Tus_far | Tussilago farfara |
| Ulm_min | Ulmus minor |
| Urt_dio | Urtica dioica |
| Val_off | Valeriana officinalis agg |
| Ver_lyc | Verbascum lychnitis |
| Ver_nig | Verbascum nigrum |
| Ver_pul | Verbascum pulverulentum |
| Ver_tha | Verbascum thapsus |
| Ver_arv | Veronica arvensis |
| Ver_cha | Veronica chamaedrys |
| Ver_off | Veronica officinalis |
| Ver_pra | Veronica praecox |
| Vib_lan | Viburnum lantana |
| Vib_opu | Viburnum opulus |
| Vic_cra | Vicia cracca agg |
| Vic_hir | Vicia hirsuta |
| Vic_sep | Vicia sepium |
| Vic_spe | Vicia spec |
| Vic_ten | Vicia tenuifolia |
| Vic_tet | Vicia tetrasperma |
| Vin_hir | Vincetoxicum hirundinaria |
| Vio_can | Viola canina |
| Vio_hir | Viola hirta |
| Vio_mir | Viola mirabilis |
| Vio_riv | Viola riviniana |
| Vio_rup | Viola rupestris |

149
